# Can a woody encroaching species affect woodlands? Impacts of eastern redcedar (*Juniperus virginiana*) overdominance on woodland understories and management scenarios

**DOI:** 10.64898/2026.11.19.767687

**Authors:** Lillian Hoffart, Bailey H. McNichol, Abigail Ridder, Brittni McGuire, Sabrina E. Russo

## Abstract

Eastern redcedar (*Juniperus virginiana* L., Cupressaceae; ERC) is rapidly expanding throughout the Great Plains. However, its effects on woodlands, particularly in the Northern Great Plains, are poorly understood, even as managers invest substantially in its removal. To fill this gap, along an ERC dominance gradient in a forest inventory plot in Nebraska, we assessed: (1) how woodland understory plant communities and environmental conditions varied and (2) where and how ERC management had the greatest benefits. Spatial neighborhood analyses showed that greater ERC canopy reduced woody seedling abundance, species richness, and species diversity relative other canopy species. No canopy species, including ERC, was associated with variation in herbaceous species richness. ERC dominance was unrelated to soil moisture, pH, or organic matter, but was associated with reduced understory light, suggesting that dense ERC foliage limits seedling establishment. Simulated management scenarios predicted that continued encroachment would shift the woody understory community toward disturbance-tolerant shrubs and away from canopy tree species, likely altering forest structural and species diversity and reducing habitat quality. ERC removal at low to moderate encroachment levels and from steep canyon habitats was associated with the greatest increases in seedling abundance and species richness, but also small declines in diversity, with implications for the complex decisions of when, where, and how much of this native species to remove. Our results indicate that ERC encroachment has reduced woody species diversity, and left unchecked, will likely continue to do so, with potentially cascading effects on higher trophic levels, biotic interactions, and ecosystem services.

## 1. Introduction

Woody plant encroachment — native woody species increasing in density and cover over time — is transforming ecosystems across the globe (S. R. Archer, 2009; Ding et al., 2020; Saintilan & Rogers, 2015). Among the most pervasive woody encroaching species in the North American Great Plains is eastern redcedar (*Juniperus virginiana* L.; Cupressaceae, hereafter “ERC”), a native conifer that was historically less common in the Great Plains but has increased dramatically in abundance and distribution in recent decades and is anticipated to continue expanding (An Atlas of the Sand Hills; Resource Atlas 5b, 1998; Engle et al., 2006; Hanberry, 2022; Twidwell et al., 2021; Yang et al., 2024). Its growing prevalence is attributed to anthropogenic factors, including greater seed availability due to shelterbelt plantings as part of the Prairie States Forestry Program — a response to the dust bowl (Dahl, 1940; Munns & Stoeckeler, 1946; Van Auken, 2008), fire suppression tied to European settlement (DeSantis et al., 2011; Donovan et al., 2018), and changes in climate (Bonannella et al., 2023; Bond & Midgley, 2000; Ratajczak et al., 2012). The effects of ERC encroachment on grasslands have been well-studied and have shown negative consequences for bird and plant diversity (Chapman et al., 2004; Limb et al., 2010, 2014), livestock production (Domeier et al., 2025), and wildfire control (Donovan et al., 2020). However, ERC has also encroached into wooded ecosystems (Meneguzzo & Liknes, 2015; Nebraska Forest Service, 2016), increasing dramatically in abundance, but its effects there are far less understood compared to effects in grasslands. Since land managers invest substantial resources into removing ERC from woodlands (Weir et al., 2017), it is important to have a more thorough understanding of how ERC affects wooded ecosystems. Therefore, the goals of this study were to quantify the effects of ERC encroachment in a Northern Great Plains woodland and to use those empirical findings to predict outcomes for woodland diversity and composition under different possible ERC management scenarios.

Several mechanisms may link ERC encroachment to cascading changes in plant communities. ERC outcompetes grassland species by dramatically reducing light availability in grasslands (Limb et al., 2010). Additionally, ERC is drought-tolerant, even while using more water than surrounding grassland species (Bahari et al., 1985; Eggemeyer et al., 2009; Kishawi et al., 2022; Zhang et al., 2026), which would allow ERC to outcompete other plants in a semiarid environment. It has also been hypothesized that ERC litter may raise soil pH due to its high calcium content (Arend 1950), which could affect soil and plant communities. A grassland study evaluating this hypothesis found small increases in soil alkalinity under ERC canopies (Broadfoot, 1951), whereas other studies found no consistent changes in soil pH (Pierce & Reich, 2010; Sauer et al., 2023; Williams et al., 2013). ERC litter is also recalcitrant to decomposition and accumulates faster compared to litter from hardwood tree and herbaceous plant species, potentially altering soil organic matter concentration, C storage, and soil microbial communities (Norris et al., 2001; Sauer et al., 2023; Williams et al., 2013). However, most of these findings are from grasslands and may not translate to wooded ecosystems, which differ in their soil properties (Hillel, 2005), light and hydrological environments, vegetation structure, and plant species composition. Whether ERC causes similar alterations in wooded ecosystems remains largely untested in the Northern Great Plains.

ERC is thought to contribute to habitat degradation of threatened oak (*Quercus* spp.) woodlands by reducing light availability in the understory which, in turn, limits the recruitment of light-demanding oak seedlings (Dey et al., 2014; Schneider et al., 2011). Much of our knowledge of encroachment by *Juniperus* species (including ERC) in woody systems comes from two sites in the US Southern Great Plains: the Cross Timbers, an oak savanna in central Oklahoma, and the Edwards Plateau, a mosaic of oak, juniper, and grassland in Texas. In the Cross Timbers, proximity to ERC trees is associated with decreased woody and herbaceous vegetation cover and species richness, largely driven by leaf litter accumulation and lack of understory light (van Els et al., 2010). Greater ERC basal area was also associated with decreased juvenile oak recruitment despite mature oaks being common throughout the study area (van Els et al., 2010), potentially signaling a transition away from oak-dominated woodland communities (Hoff et al., 2018). In the Edwards Plateau, ashe juniper (*Juniperus ashei*), a close relative of ERC, is growing in increasing numbers across the oak-juniper woodlands and leading to the “juniperization” of otherwise more diverse forests (Fowler & Carden, 2024). Increasing proximity to ashe juniper resulted in decreased herbaceous cover and species richness, which were associated with juniper-driven changes in microenvironment, including canopy closure, producing more uniformly dark understories with reduced light niche variation (Fuhlendorf et al., 1997; Yager & Smeins, 1999).

ERC encroachment rates are accelerating (Shrestha et al., 2025), and ERC is increasingly dominating woodland systems within its current range (Meneguzzo & Liknes, 2015; Nebraska Forest Service, 2016). Land managers seeking to conserve threatened habitats like oak woodlands and preserve biodiversity and ecosystem function of woodlands spend considerable resources removing ERC. The cost of one treatment plan for restoring ERC-encroached oak-hickory, oak-pine, and post oak-blackjack oak forest approximated $180,000 for 5,000 acres for only an initial treatment (Weir et al., 2017). In response to the monumental amount of encroachment facing managers, the “defend the core” approach has been gaining traction (Fogarty & Twidwell, 2023; Twidwell et al., 2021). The principle of “defend the core” is to concentrate management in areas with little-to-no encroachment in intact, un-encroached areas and surrounding areas with light to moderate encroachment, rather than focusing on areas with the most severe encroachment. While this approach makes sense intuitively, it has not been explicitly evaluated in the context of woodland management.

In order to understand the conservation implications of ERC encroachment in Northern Great Plains woodlands, we investigated how ERC dominance in the overstory impacts woodland understory community structure (abundance, species richness, diversity, and composition of woody seedlings; cover, species richness, and composition of herbaceous plants) and environmental conditions (soil pH, moisture, and organic matter concentration and understory light availability) in a 20.2 ha forest inventory plot located on the south bank of the Niobrara River, Nebraska, USA. Classified as a Biologically Unique Landscape by Nebraska Game and Parks (Schneider et al., 2011), the Niobrara woodlands lining the steep slopes of the spring-fed canyons are of great conservation importance, being a post-glacial climatic refugium and migration corridor (Bessey, 1887; Kantak & Churchill, 1993; Kaul et al., 1988). Supporting plant species from eastern deciduous, western coniferous Rocky Mountain, and northern boreal regions, many of the species found in these woodlands are at the extremes of their geographic ranges, potentially leading them to be vulnerable to stress and disturbance, including competition from encroaching species (Robertson et al., 2019). However, these woodlands are threatened by increased tree densities, primarily of ERC (Schneider et al., 2011). ERC, while native to Nebraska, was not historically found in large numbers in these canyons until recently (An Atlas of the Sand Hills; Resource Atlas 5b, 1998).

We tested the hypothesis that, as a woody encroaching species, if ERC is transforming understory community structure in ways that reduce woodland diversity and shift species composition, then we would expect ERC’s effects to be stronger than those of other dominant canopy tree species that are not considered to be woody encroaching species at this site: bur oak (*Quercus macrocarpa* Michx.), American basswood (*Tilia americana* L.), green ash (*Fraxinus pennsylvanica* Marsh.), and common hackberry (*Celtis occidentalis* L.). Moreover, examining the effects of ERC on understory environmental conditions (soil moisture, soil pH, organic matter concentration, and understory light availability) may provide insights into possible mechanisms by which ERC encroachment may alter understory communities. After identifying response variables that were negatively affected by increasing ERC overstory dominance, we used our empirical models to simulate ERC thinning to make predictions about how these variables would respond under different ERC management scenarios ranging from no management (continued ERC encroachment) to high levels of management (ERC removal) and about the topographic habitats in which management would have the greatest conservation effect. We addressed the following research questions:

*Q1: How does eastern redcedar affect the woody plant community in the understory compared to other dominant canopy species?* If ERC limits the recruitment of other woody species in the understory, then greater ERC dominance will reduce the abundance, species richness, diversity, and evenness of woody species and result in shifts in woody species composition compared to other canopy tree species.

*Q2: How does eastern redcedar affect the herbaceous plant community in the understory compared to other dominant canopy species?* If ERC limits herbaceous plant recruitment in the forest understory, then greater ERC dominance will reduce the cover and species richness of herbaceous plants and result in shifts in herbaceous species composition compared to other canopy tree species.

*Q3: How does eastern redcedar affect understory environmental conditions compared to other dominant canopy species?* If ERC’s alteration of the understory environment is a mechanism for changes in plant communities, then environmental conditions where ERC is more dominant will differ from areas with greater dominance of other canopy species. Specifically, based on previous findings described above, environmental conditions with ERC dominance will be characterized by drier, more alkaline soils, less soil organic matter, and less understory light.

*Q4: What are the conservation implications of different eastern redcedar management scenarios?* Using the empirical models from research questions 1-3, we (1) predicted how the understory would respond under simulated thinning scenarios, from no management (increasing ERC dominance and replacement of canopy species) to management by varying amounts of ERC removal, (2) evaluated whether management would be more effective in particular topographic habitats, and (3) evaluated whether management effectiveness depended on the initial level of ERC encroachment. We predicted that the understory response to ERC removal would depend on topography. We also predicted that removal would benefit woodland diversity most at lower levels of initial ERC encroachment, consistent with the “defend the core” management strategy (Twidwell et al., 2021).

## 2. Methods

### 2.1. Study site

This study was conducted at the Niobrara Forest Plot (42.780° N, 100.021° W), located at The Nature Conservancy’s Niobrara Valley Preserve in Brown County, Nebraska. The 20.2 ha (560 m × 360 m) plot was established in 2019 and is part of the Smithsonian Forest Global Earth Observatory network (Davies et al., 2021). The region has a semiarid climate (2000 – 2025 mean annual temperature of 9.5 °C, average daily minimum and maximum temperatures of 2.9 and 16.2 °C, mean annual total precipitation of 57 cm; PRISM Group; Oregon State University, accessed May 2026). The Niobrara Plot is in a woodland bordered by the Niobrara River to the North and by sandhills grassland to the south, creating a dramatic transition from forest to prairie along a 59-meter elevation gradient (McNichol et al., 2024). Soils are well-drained and derived from Holocene-aged eolian sand deposits, which allow for rapid precipitation infiltration with little to no runoff (Bleed & Flowerday, 1998). Seeps and springs are common in canyons throughout the woodlands in this region and drain to the Niobrara River (Hearty, 1978; Kaul et al., 1988). ERC management in the Niobrara Plot consisted of thinning in 2017 with machinery and manual hand tools, followed by a prescribed burn in 2020. In 2019, ERC made up 34% of stems in the Plot (Table 1), remaining the most abundant woody species in the woodland despite management.

**Table 1:**
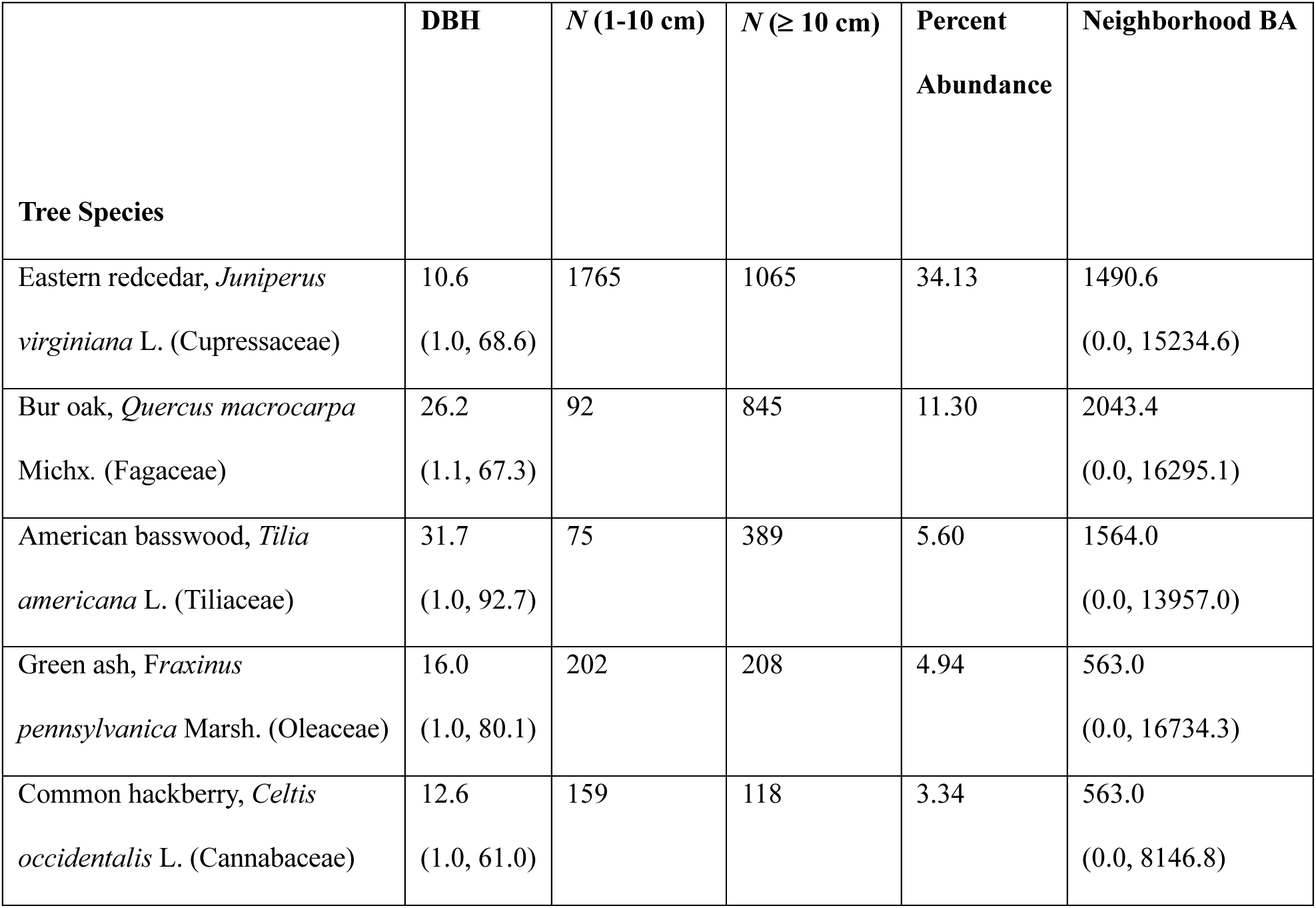
Abundance, diameter, distribution, and neighborhood dominance of the five most abundant canopy tree species in the Niobrara Plot. For each species, the table shows the average tree diameter at breast height (DBH, cm) and DBH range in parentheses, number (*N*) of trees 1-10 cm DBH and ≥10 cm DBH, and the percent abundance of all trees in the Niobrara Plot, as well as the mean basal area (BA; cm^2^) with the range in parentheses of all stems in 15-meter neighborhoods (see Figure S1)

| **Tree Species** | **DBH** | ***N* (1-10 cm)** | ***N* (≥ 10 cm)** | **Percent Abundance** | **Neighborhood BA** |
|---|---|---|---|---|---|
| Eastern redcedar, *Juniperus virginiana* L. (Cupressaceae) | 10.6 (1.0, 68.6) | 1765 | 1065 | 34.13 | 1490.6 (0.0, 15234.6) |
| Bur oak, *Quercus macrocarpa* Michx. (Fagaceae) | 26.2 (1.1, 67.3) | 92 | 845 | 11.30 | 2043.4 (0.0, 16295.1) |
| American basswood, *Tilia americana* L. (Tiliaceae) | 31.7 (1.0, 92.7) | 75 | 389 | 5.60 | 1564.0 (0.0, 13957.0) |
| Green ash, F*raxinus pennsylvanica* Marsh. (Oleaceae) | 16.0 (1.0, 80.1) | 202 | 208 | 4.94 | 563.0 (0.0, 16734.3) |
| Common hackberry, *Celtis occidentalis* L. (Cannabaceae) | 12.6 (1.0, 61.0) | 159 | 118 | 3.34 | 563.0 (0.0, 8146.8) |

### 2.2. Data collection

#### 2.2.1. Tree inventory data

The Niobrara Plot is gridded into 504 20 m^2^ quadrats with all corner intersections marked with a georeferenced survey pole. In 2019, all woody stems ≥ 1 cm in diameter at breast height (∼1.3 m; DBH) were tagged, identified to species, and measured for DBH, for a total of 8,294 trees across 27 species (McNichol et al., 2024; Russo, 2026). We mapped each tree’s location to the nearest ∼20 cm, but inventory data were collected at the stem level because many individual trees possess multiple stems due to resprouting. Understory plant community composition and environmental variables (see below) were collected in subsequent years from a 1 m^2^ subplot (Table S1) near the northeast corner of each quadrat. We used the tree inventory data to calculate the dominance of each of the five most abundant canopy tree species in a neighborhood surrounding each subplot (Figure 1; described below).

**Figure 1:**
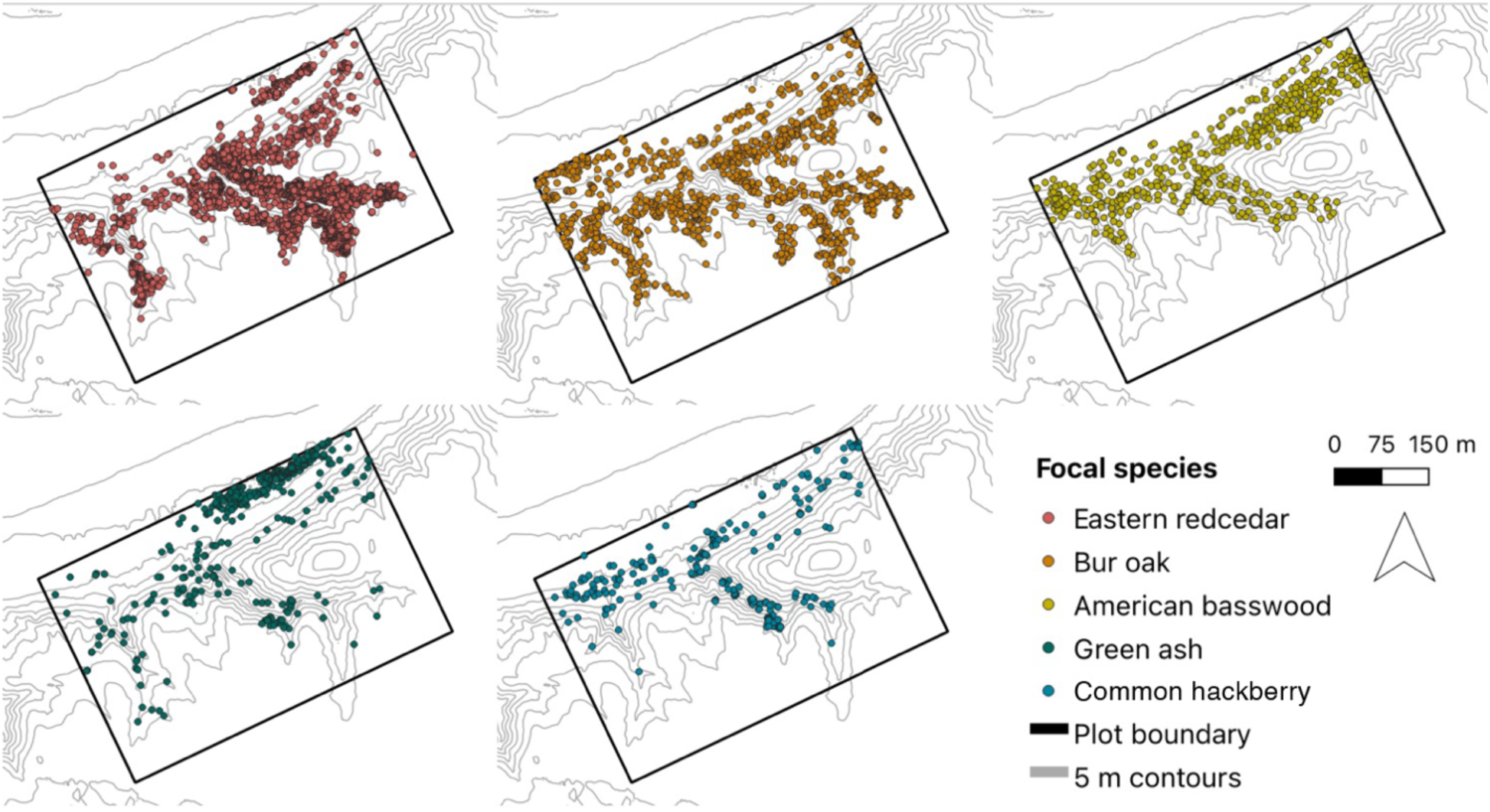
Distribution of focal canopy tree species in the Niobrara Plot, Nebraska. Dots represent trees with diameter at breast height ≥ 1 cm, the Plot boundary is outlined in black, and 5-m elevation contours are shown with gray lines.

#### 2.2.2. Topographic variation

Using a 1-m resolution digital elevation model (DEM) derived from remotely sensed light detection and ranging (LiDAR) data (U.S. Geological Survey, 2023), topographic information (elevation, m; slope, %; and aspect decomposed into northness and eastness) was extracted for each subplot. Northness and eastness were calculated as the sine of the slope, multiplied by the cosine and sine of the aspect, respectively (Amatulli et al., 2018).

#### 2.2.3. Understory plant community

In the summers of 2021 and 2022, we censused woody plants < 1 cm DBH at each of the 504 1-m^2^ subplots. Woody plants included tree seedlings with adult growth forms that exceed 1 cm in DBH and other species such as shrubs that never exceed 1 cm in DBH; these are collectively referred to here as “seedlings,” even though some of these plants were not juveniles. We also conducted a groundcover survey in 2021 in all subplots and classified the percent cover of herbaceous plants, woody plants, bare ground, and woody debris in each subplot. Percent cover of vegetation was calculated by summing herbaceous and woody plant percent cover (McNichol & Russo, 2026). In July and August of 2021, we surveyed herbaceous vegetation in 78 subplots by recording the presence or absence of all non-woody plant species (sedges, forbs, grasses, non-vascular plants, and lower vascular plants) (McNichol & Russo, 2026). For the two cases in which species-level identification was not possible, morphospecies designations were used. The 78 subplots were chosen in a stratified random design to capture the full range of topographic habitats within the Niobrara Plot (McNichol et al. 2024).

#### 2.2.4. Environmental variables

To estimate understory light availability, we measured canopy openness (%) in July 2022 at all 504 subplots using a concave spherical densiometer. At 68 of those subplots, we also took hemispherical photographs during uniform overcast sky conditions of the canopy ∼1 m above the soil using a Nikon CoolPix 4500 camera with an FC-E8 fisheye lens converter. The photographs were analyzed using WinSCANOPY software (v. 2008a, Regent Instruments, Canada) to estimate canopy openness (%) and total daily photosynthetically active photon flux density (PPFD, mol/m^2^/day), averaged over the year for each subplot. To obtain PPFD estimates for all subplots, we used a linear model to predict PPFD from canopy openness based on spherical densiometer measurements for subplots where both were quantified (PPFD = 0.5013 × Canopy Openness + 4.5375; *R*² = 0.86) and used that model to predict PPFD for all subplots based on the densiometer-based Canopy Openness measurements.

At 74 subplots chosen to capture the full range of topographic habitats, we measured soil volumetric water content (VWC, %) using a handheld time-domain reflectometer (Campbell Scientific Hydrosense II) from May to October 2021. On a single, rain-free day each month, three replicate measurements were taken at each subplot. These were averaged first by subplot, then averaged across months, to obtain a mean soil moisture value per subplot (McNichol et al., 2024). In August 2022, a soil core (5 cm diameter x 10 cm depth) was collected next to 50 subplots and analyzed for soil texture (particle size analysis; Robertson et al., 1999), soil pH (Sparks et al., 1996), and soil organic matter (loss on ignition, % LOI; Schulte & Hopkins, 1996; Combs & Nathan, 1998).

### 2.3. Statistical analysis

We estimated the effect of five focal canopy species’ dominance (Table S1) on understory communities and environmental conditions using neighborhood models fit with Bayesian regression, implemented in Stan (Carpenter et al., 2017) via the *brms* package (Bürkner, 2017) in R statistical software (version 4.1; Posit team, 2024). All models shared a common set of fixed-effect predictors — neighborhood dominance of each focal species, topographic position (topo-PC1 and topo-PC2) and understory light (PPFD) — for each response variable: woody seedling abundance, richness, evenness, and Shannon diversity; vegetative groundcover; herbaceous species richness; soil VWC, pH, and organic matter; and understory light (Table S1). To account for annual variation, year was included as a fixed-effect predictor for response variables in which seedling data had been collected in two years. All continuous predictors were standardized with z-score standardization. Understory light was not included as a covariate in the model for which PPFD was the response variable.

#### 2.3.1. Neighborhood dominance

In a radius around each subplot (neighborhood), we calculated neighborhood dominance of each species using the stem-level Plot data and the location of each tree to determine which trees were in each neighborhood. We used the 371 subplots for which there was ≥ 1 tree (DBH ≥ 1 cm) in the neighborhood and compared three different candidate neighborhood metrics and two radii (10 and 15 m) to identify the one that best captured the effects of the overstory. These radii were selected as they are approximately the size of a crown of a large canopy tree crown at this site. The candidate metrics were: (1) Stem density, or the proportion of stems comprised of the focal species in a neighborhood, *N/T*, where *N* is the number of stems of the focal species, and *T* is the number of stems of all species. (2) Basal area, or the proportion of stem basal area comprised of the focal species in a neighborhood, 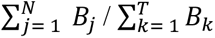, where *B* is the basal area of a stem calculated as *π* * (DBH/2)^2^ with *j* indicating the focal species, and *k* indicating all species. (3) Proportional basal area with distance-based exponential decay, or the proportion of stem basal area comprised by the focal species in a neighborhood, weighting stems nearer the center of the neighborhood more based on an exponential decay, 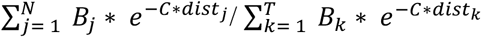, where *dist* is the distance from each stem to the center of the neighborhood and *C* is the distance decay constant log(2) / 0.5 *R,* with *R* representing the radius of a stem (Uriarte et al., 2004; Zhao et al., 2006).

We used leave-one-out cross-validation (LOO-CV; Vehtari et al., 2017) to compare models using the different metrics and radii. The most supported neighborhood metric was based on the difference in the expected log predictive density (ELPD), where the model with the lowest ELPD value was selected. Models that fell within 2 standard errors (SE) of the top model were considered equivalent, and thus the simplest model that captured data structure was selected. For most response variables, the selected model was the one containing the proportion of basal area of the focal species (metric number two above) in a 15 m radius, which was used in all proceeding analyses. For the neighborhood dominance values for all species ranged from 0, representing no trees of that species in the neighborhood, to 1, representing a neighborhood entirely composed of that species (Figure S1).

#### 2.3.2. Topography variables

Because topography affects plant community composition and overstory basal area (McNichol et al. 2024), we included topography as a covariate in all models. We used principal components analysis (PCA) to derive orthogonal axes summarizing the topographic position (elevation, slope, eastness, and northness) of each subplot. Principal components (PC) 1 and 2 accounted for 73.4% and 23.4% of variance, respectively (topo-PC1, topo-PC2; Table S2).

#### 2.3.3. Q1: How does eastern redcedar affect the woody plant community in the understory compared to other dominant canopy species?

We modeled the effect of tree species dominance on the understory woody plant community (woody seedling abundance, species richness, evenness, and Shannon diversity). All predictor variables were scaled prior to modeling. For likelihood families, we used a negative binomial distribution for abundance to model overdispersed count data, a Poisson distribution for species richness to model count data, a beta distribution for evenness, transformed following Smithson & Verkuilen, 2006 in order to fit the assumption of a beta distribution (0 < *x* < 1), and a hurdle-gamma distribution for Shannon diversity to model continuous data with excess zeros. For the richness model, we only assessed neighborhoods with richness ≥ 1.

For each response variable, we performed model selection with three candidate models with increasing levels of complexity using LOO-CV in the *loo* package (Vehtari et al., 2017). The three candidate models differed in how each canopy species impact interacted with topographic position (topo-PC1 and topo-PC2). Model 1 included all five species’ dominance, topo-PC1, topo-PC2, as main effects only with no interaction terms. Model 2 was an extension of model 1, adding two-way interactions between each species’ dominance and topo-PC1. Model 3 extended model 2 by crossing each species’ dominance term with both topographic variables (species’ dominance × topo-PC1 × topo-PC2), which includes all two-way interactions in addition to three-way interactions. PPFD and year were included as covariates in all three models and did not interact with species or topographic terms. Model selection followed the steps described section 2.3.1 (Table S3). We evaluated multicollinearity of non-interactive predictors in the final models using variance inflation factors (VIF); predictors with VIF > 5 were removed as evidence of multicollinearity (two cases: topo-PC2 in the soil pH and organic matter models).

Because the species effects on the Shannon diversity model appeared non-linear in the species effects plots, we additionally tested quadratic terms for species dominance. Using LOO-CV and the evaluation criteria described above, we compared combinations quadratic and linear terms for species dominance. The final Shannon diversity model included a quadratic term for bur oak, ERC, and green ash dominance.

We tested our models for evidence of spatial autocorrelation using Moran’s *I* (Legendre, 1993; Moran, 1950) in the *ape* package (Paradis & Schliep, 2019). We considered the model results to be affected by spatial autocorrelation if Moran’s *I* was significant (*p* ≤ 0.05). Three models — woody seedling abundance, richness, and diversity — exhibited statistically significant values. We were unable to fit CAR terms in the diversity model due to lack of convergence, but for seedling abundance and richness, we incorporated a conditional autoregressive (CAR) term (Ver Hoef et al., 2018). Adding the CAR term significantly reduced the effect of spatial autocorrelation in all cases (Table S4).

We specified weakly informative priors to allow the data to dominate inference in all models. We used a normally distributed prior with a mean of 0 and standard deviation of 10 for fixed effects (slope and intercept parameters) for all models. For priors that included a CAR term, we used an exponentially distributed prior with a rate parameter of 1 for the standard deviation. For all other terms, we retained the default brms priors (flat, non-informative; Bürkner, 2017). To verify that our results were not sensitive to our choices of prior, we conducted a prior sensitivity analysis by doubling and halving (e.g. Normal (0, 20), Normal (0, 5)) the standard deviation of the fixed-effect priors and refitting the models. Models were considered robust if model coefficients and 95% credible intervals were similar across all prior specifications (Banner et al., 2020); we found no dependence of coefficients on prior specification. We also verified that our choice of priors produced biologically plausible estimates with prior predictive checks (Lemoine, 2019).

We assessed model convergence by visually inspecting model trace plots, posterior predictive checks, and ensuring the Gelman-Rubin *R̂* statistic was < 1.05 for each parameter (Gelman et al., 2012). Non-CAR models were run with four chains of 2,000 warmup and 4,000 total iterations whereas models including CAR terms required four chains of 8,000 warmup and 16,000 total iterations to converge. We computed the Pareto shape parameter, *k*, which can be used to identify overinfluential observations (Vehtari et al., 2017); over 95% of *k* values were < 0.7 in all models.

Mean parameter estimates and 95% credible intervals were calculated from the posterior distribution of each parameter. Because the bur oak, ERC, and green ash dominance parameter in the Shannon diversity model contained quadratic terms, we calculated the mean parameter estimated and credible interval at the mean value of each species’ dominance (ERC = 0.18, bur oak = 0.31, green ash = 0.08). Our hypothesis compares the influence of ERC with other focal tree species on our response variables. We therefore generated *a-priori* planned contrasts of the parameter estimates for ERC versus the other four focal species by subtracting posterior draws for each tree species’ slope parameters from the corresponding draw for ERC. Contrasts with a 95% credible interval excluding zero were interpreted as evidence that the effects of ERC were different than those of other focal species. Unlike other modeling approaches, Bayesian models avoid the need for multiple comparison corrections when estimating contrasts, as these comparisons are functions of the same joint posterior distribution rather than independent tests (Berry & Hochberg, 1999). Additionally, comparisons were specified *a priori* in planned contrasts, avoiding data-driven additional tests (Gelman & Loken, 2013).

To evaluate the effects of species dominance and environmental variables on understory woody seedling species composition, we performed distance-based redundancy analysis (dbRDA; *vegan* package; Oksanen et al., 2026) using the dominance of the five focal species and environmental variables (topo-PC1 and PC2, PPFD) as constraining variables based on Bray-Curtis distances (Legendre & Andersson, 1999). We removed four subplots that had extreme and unrepresentative woody species composition (many individuals of one species found nowhere else in the Niobrara Plot). Species relative abundance was transformed using a Hellinger transformation, as many species were only represented in a few subplots (Legendre & Gallagher, 2001). Overall model significance and significance of individual constraining variables were tested using analysis of variance (*anova.cca; vegan)*, and adjusted *R*^2^ values (*vegan;* Legendre et al., 2011).

#### 2.3.4. Q2: How does eastern redcedar affect the herbaceous plant community in the understory compared to other dominant canopy species

Estimations of ERC influence on predictors related to herbaceous plant community (vegetative groundcover, herbaceous species richness) were modeled using the workflow described in section 2.3.3. Herbaceous data were collected in one year, so year was not included in the model as a covariate. For likelihood families, we used a beta distribution for vegetative groundcover which fit our data (0 < *x* < 1) and a Poisson distribution for herbaceous species richness to model count data. Moran’s *I* tests indicated that the vegetative groundcover model exhibited significant spatial autocorrelation (Table S4), so a CAR term was included in the model, which resulted in non-significant spatial autocorrelation.

#### 2.3.5. Q3: How does eastern redcedar affect understory environmental conditions compared to other dominant canopy species?

Estimations of ERC influence on environmental conditions (soil VWC, pH, and organic matter; PPFD) were modeled using the workflow described above for Q1. Environmental data were collected in one year, so year was not included in the model as a covariate. For likelihood families, we used a lognormal distribution for soil VWC, as these data were overdispersed and continuous, and normal distributions for soil pH and soil organic matter. Soil pH values were centered around 0 prior to modeling. For the PPFD model, to achieve model convergence, the predictor variables of bur oak, American basswood, green ash, and common hackberry dominance were grouped into a single predictor variable representing non-ERC canopy cover. We standardized PPFD measurement on a 0-1 scale and applied the Smithson & Verkuilen (2006) transformation to accommodate values at 0 and 1, modeling PPFD with a mixture of two beta distributions to capture its bimodality, representing high and low light conditions. Moran’s *I* tests indicated that the VWC and PPFD models exhibited significant spatial autocorrelation, but we were unable to fit CAR terms due to lack of convergence (Table S4). Priors, model convergence, parameter estimates, and 95% credible intervals were examined following the steps section 2..

#### 2.3.6. Q4: What are the conservation implications of different eastern redcedar management scenarios?

To address Q4, we simulated ERC management scenarios from none to varying levels of ERC removal (as a range of the percent of ERC basal area removed) by adjusting the amount of ERC, other canopy trees, and light in our observed dataset and then using the empirical models from Questions 1-3 to predict how the understory would respond. We focused on the understory variables significantly affected by ERC, namely woody seedling abundance, species richness, Shannon diversity, and community composition (see *Results*). While this approach of estimating the effects of management on woodland communities has the disadvantage of extrapolating the model beyond the observed data, it has numerous advantages. It generates predicted conservation outcomes under contrasting management scenarios using empirical results. This provides managers with an evidence-based approach to selecting a management strategy without the considerable expense of time (including waiting for systems to respond) and resources required to conduct an experiment to compare alternative management scenarios. Given the rapid expansion of ERC, timely management combatting encroachment and conserving diversity and ecosystem function is increasingly urgent. We also addressed the following Management Questions concerning ERC removal: (1) what initial levels of ERC encroachment should be targeted for removal in order to have the greatest increase in understory seedling abundance and richness? (2) What topographic variables are associated with the greatest increase in understory seedling abundance and richness resulting from removal? (3) Is there a predicted shift in woody species community composition between two management extremes: lack of management (continued encroachment) and nearly complete (95%) ERC removal?

To address Management Q1, we predicted the effects of removal of 35%, 55%, 75%, and 95% of the basal area of ERC in each neighborhood based on the models of woody seedling abundance, richness, and Shannon diversity described in Q1. Based on preliminary analysis, we assume ERC encroachment replaces other canopy species, so we also created a no-management scenario of continued encroachment in which ERC basal area increased by 10%, with a corresponding reduction in the basal area of other canopy tree species, evenly applied. First, we simulated ERC removal from each neighborhood in our dataset by subtracting ERC basal area from observed values and recalculating the proportion of basal area of each focal canopy species after ERC removal, which were used as new data in the prediction model.

Second, since removing canopy cover allows increased light penetration to the understory, we modeled the expected response of PPFD from the reduced total basal area. We fit a model of PPFD using the predictors of total basal area, topo-PC1, and topo-PC2, using a mixture distribution with two gaussian components as the likelihood family, using the workflow described above for Q1. A mixture distribution was selected because the two gaussian components likely each represented a different light environment occurring at high frequency in our data: low-light forest understory and high-light ecotone and canopy gaps. From the PPFD model, we obtained predicted PPFD for each subplot after ERC removal, which was used as new data in the prediction model. Using the new data, we predicted woody seedling abundance, richness, and Shannon diversity under these management scenarios and plotted the predicted change in woody seedling abundance, richness, and Shannon diversity after ERC removal along an ERC dominance pre-removal gradient. To address Management Q2, we then plotted the predicted woody seedling abundance, richness, and Shannon diversity after each ERC removal level against topo-PC1 and topo-PC2 in separate models.

To address Management Q3, we evaluated the predicted change in the understory woody seedling community composition under a lack of ERC management, resulting in increasing ERC dominance expected from worsening encroachment and from ERC management with removal of 95% of ERC basal area. We made these predictions based on our dbRDA model described in Q1. First, we simulated the effects of increasing ERC dominance by subtracting 10% of each non-ERC focal species and adding that amount of basal area to ERC, representing a change in overstory composition without changing total basal area. For each non-ERC focal species, we calculated the change in dbRDA1 and dbRDA2 between the observed and replacement scenarios. To quantify uncertainty in the predicted community shifts, we generated 1,000 bootstrap resamples of the predicted communities per scenario and constructed 95% confidence ellipses for each focal species. To identify the woody seedling species most strongly associated with compositional change following ERC replacement, species scores were calculated as weighted averages of species abundances using the wascores function in the *vegan* package. Species with the greatest scores along each ΔdbRDA axis, corresponding to community shifts, were interpreted as changing the most. We tested if the predicted species shifts were significant and directional with Rayleigh tests, which are used to examine the unimodal concentration of vectors (Watanabe, 2022). Then, we used a similar workflow to predict the change in community composition in each neighborhood under the 95% ERC removal scenario from the dbRDA model.

## 3. Results

### 3.1. Q1: Eastern redcedar affected the understory woody plant community more strongly than other canopy species

As the dominance of ERC in the canopy increased, woody seedling abundance decreased sharply (slope = −1.693, 95% CI [-2.598, −0.804], Table S5, Figure 2A; Figure S2), whereas there was no effect of any other focal species on woody seedling abundance (Table S5, Figure 2A). Moreover, the effect of ERC canopy dominance on woody seedling abundance was substantially more negative than most other codominant tree species, as seen from the mean difference in slope estimates relative to ERC (bur oak: 0.982, 95% CI [0.138, 1.821]; American basswood: 1.981, 95% CI [0.882, 3.078]; green ash: 0.554, 95% CI [-0.645, 1.740]; common hackberry: 1.843, 95% CI [0.134, 3.530], Table S6). Likewise, ERC was the only species for which increasing dominance reduced woody seedling species richness (slope = −0.655, 95% CI [-1.063, −0.250], Table S5, Figure 2B, Figure S3), whereas there was no effect of increased bur oak, American basswood, green ash, or common hackberry dominance on woody species richness (Table S5, Figure 2B). The effect of ERC dominance on woody seedling richness was more strongly negative than that of bur oak and American basswood (bur oak: 0.688, 95% CI [0.286, 1.091], American basswood: 0.992, 95% CI [0.534, 1.445], and green ash: 0.695, 95% CI [0.1667, 1.221], Table S6). Woody seedling evenness did not vary strongly with dominance for any of the canopy species tested (Table S5, Figure 2C, Figure S4), and the effect of ERC dominance was not different from any other focal species (Table S6). Woody seedling Shannon diversity was negatively impacted by ERC and common hackberry (ERC: slope = −1.220, 95% CI [-1.979, −0.439]; common hackberry: slope = −0.701, 95% CI [-1.249, −0.138]; Table S5, Figure 2D, Figure S5). ERC dominance had a more strongly negative effect on Shannon diversity than all canopy species except hackberry (0.519, 95% CI [-0.428, 1.478]; Table S6).

**Figure 2:**
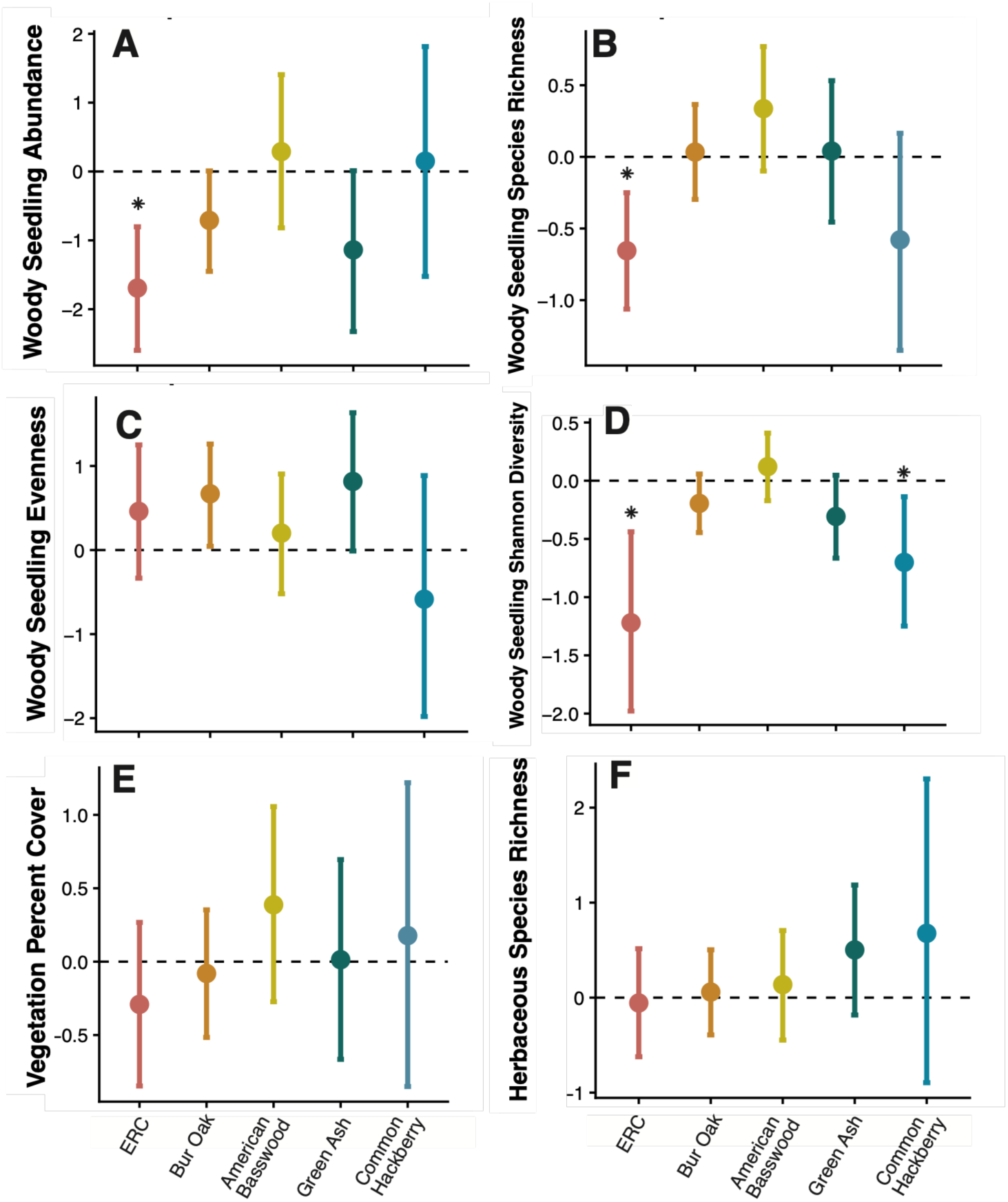
Effects of canopy species on woody and herbaceous understory communities in the Niobrara Plot, Nebraska. Slope parameter estimates and their 95% credible intervals are shown for woody seedling (A) abundance, (B) species richness, (C) evenness, and (D) Shannon diversity, where, due to a quadratic term in the model, ERC, bur oak, and green ash slopes were evaluated at a dominance of 0.50, (E) understory vegetation percent cover, and (F) herbaceous species richness. Stars indicate credible intervals that exclude zero. ERC = eastern redcedar; oak = bur oak; basswood = American basswood; ash = green ash; hackberry = common hackberry.

Species composition of the understory woody plant community was significantly affected by canopy species dominance, topography, and light (*F* = 7.961, *p* = 0.001, *adjusted R*² = 0.163), with dbRDA axis one and two accounting for 14.0% and 1.7%, respectively, of variation in seedling community composition. The dominance of ERC accounted for a significant but small percentage of the variation in species composition of the woody understory community, and all species except green ash were also significantly associated with variation in woody understory composition (ERC: *F* = 1.812, *p* = 0.047; bur oak: *F* = 2.401, *p* = 0.002; American basswood: *F* = 2.796, *p* = 0.003; common hackberry: *F* = 2.015, *p* = 0.017, Figure S6A).

### 3.2. Q2: Eastern redcedar did not affect the understory herbaceous plant community more strongly than other canopy species

We found that percent cover of understory vegetation was not affected by variation in the dominance of any canopy species, with all species exhibiting similar effect sizes to ERC (Table S5, Figure 2E, Table S6, Figure S7). Species richness of understory herbaceous plants was also not affected by any variation in the dominance of any canopy species, with all species exhibiting similar effect sizes to ERC (Table S5, Figure 2F; Table S6, Figure S8).

Species composition of the understory herbaceous community was affected by canopy species dominance, topography, and light (*F* = 3.254, *p* = 0.001, adjusted *R*² = 0.196), with dbRDA axis one and two accounting for 16.0% and 3.6%, respectively, of variation in herbaceous plant composition. However, the effect of ERC dominance on herbaceous plant composition was not significant; rather, bur oak, American basswood, and green ash had significant effects (bur oak: *F* = 1.790, *p* = 0.049; American basswood: *F* = 2.708, *p* = 0.004; green ash: *F* = 1.749, p = 0.038; Figure S6B).

### 3.3. Q3: Eastern redcedar affected understory light in low light environments more strongly than other canopy species

For none of the canopy species did variation in dominance influence soil VWC, soil pH, or soil organic matter, with all species exhibiting similar effect sizes to ERC (Table S5, Figure 3, Table S6, Figure S9, S10, S11). PPFD was affected by ERC dominance in low-light environments but not high-light environments (low light: −1.149, 95% CI [-1.779, −0.517]; high light: 0.007, 95% CI [-0.713, 0.739]; Table S5, Figure 3D; Figure S12), whereas combined dominance of the other canopy species did not affect PPFD. ERC’s impact on PPFD was significantly more negative than the other canopy species in low light environments (1.061, 95% CI [0.468, 1.654]; Table S6).

**Figure 3:**
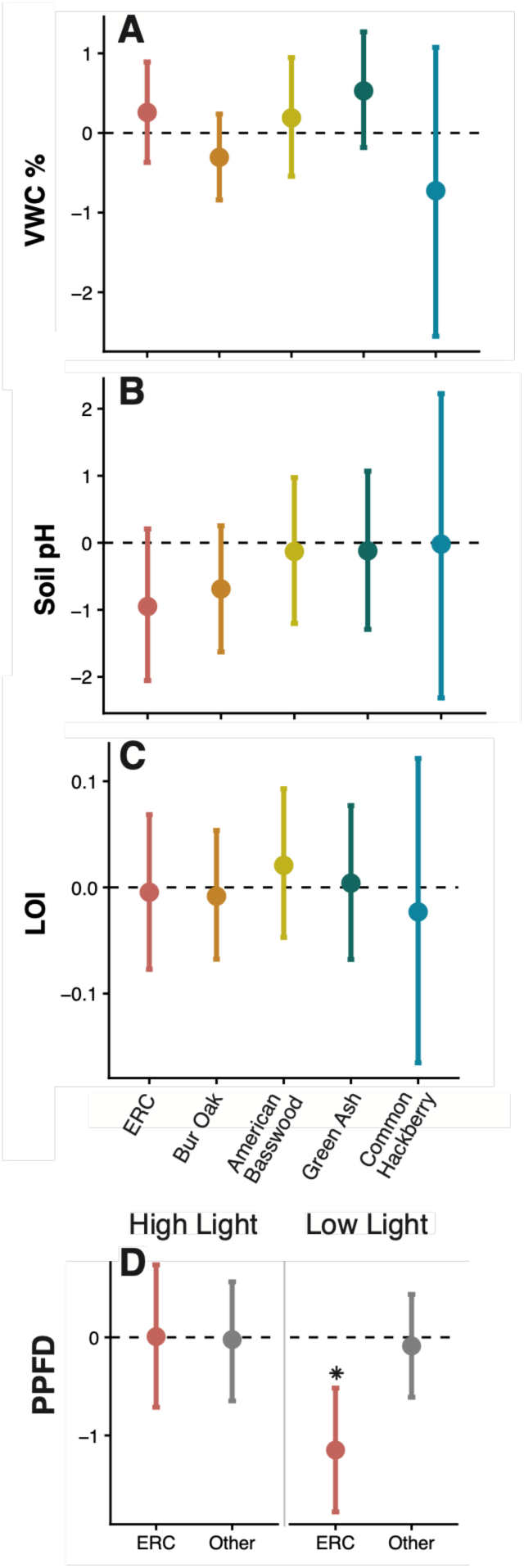
Effects of canopy species on understory environmental conditions in the Niobrara Plot, Nebraska. Slope estimates for focal species’ impacts on (A) volumetric water content (VWC, %), (B) soil pH, (C) soil loss on ignition (LOI, %), and (D) Photosynthetic Photon Flux Density (PPFD; μmol/m²/s) in high- and low-light environments. ERC = eastern redcedar; oak = bur oak; basswood = American basswood; ash = green ash; hackberry = common hackberry.

### 3.4. Q4: Simulated ERC removal predicts increases in seedling abundance and richness, particularly in forest with low to moderate densities of ERC, and lack of management predicts changes to the woody seedling community

Woody seedling abundance and species richness in the understory markedly increased in all simulated ERC removal scenarios (Figure 4). The more basal area of ERC that was removed, the greater the predicted post-removal increase in woody seedling abundance and richness. Removing ERC was predicted to increase woody seedling abundance most in absolute gains where pre-removal ERC basal area was 30-50% (Figure 4A), and most in percentage gains where it was 50-80% (Figure 4B). Woody seedling richness showed the same pattern, with the greatest absolute gains at 30-50% (Figure 4C) and the greatest percent gains at 50-80% (Figure 4D). However, removing ERC was predicted to increase woody seedling Shannon diversity the most in absolute gains when pre-removal ERC basal area is high (> 60%; Figure 4E), with the greatest absolute gains > 60% (Figure 4F). Lack of removal — continued encroachment and replacement of other canopy species — resulted in little to negative predicted change for woody seedling abundance, richness, and diversity. For the 95% removal scenario, post-removal increases in woody seedling abundance and richness were predicted to be greater for woodland topographic habitats with north- and east-facing and steeper slopes (higher values of topo-PC1) (Figure 5A, Figure 5C) and for mid-range elevations (values of topo-PC2 between −1 and 0) (Figure 5B, Figure 5D), but there was no strong relationship between seedling diversity and topography (Figure 5E, 5F).

**Figure 4:**
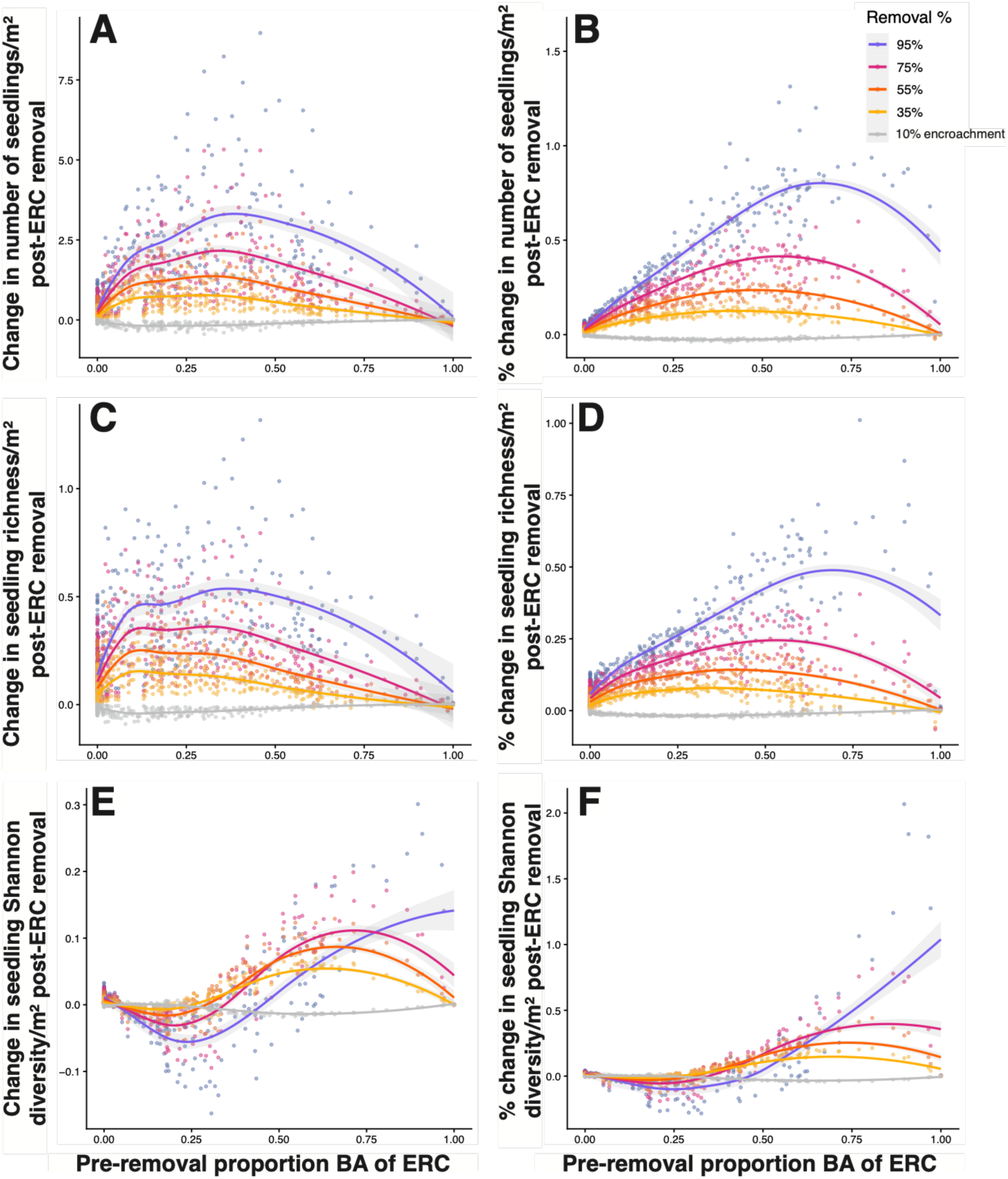
Predicted conservation effects of different simulated ERC management scenarios. Thinning scenarios were defined by 35-95% removal of ERC compared to a lack of removal, in which ERC basal area increases by 10%. Projections of the change in seedling abundance and richness across pre-removal proportion basal area of ERC following simulated ERC removal in neighborhoods. The colors represent scenerios where 95%, 75%, 55%, and 35% of ERC is removed from a neighborhood. (A) Change in seedling abundance per m², (B) percent change in seedling abundance, (C) change in seedling richness, (D) percent change in seedling richness, (E) change in seedling Shannon diversity, and (F) percent change in seedling Shannon diversity along a gradient of the pre-removal proportion BA of ERC.

**Figure 5:**
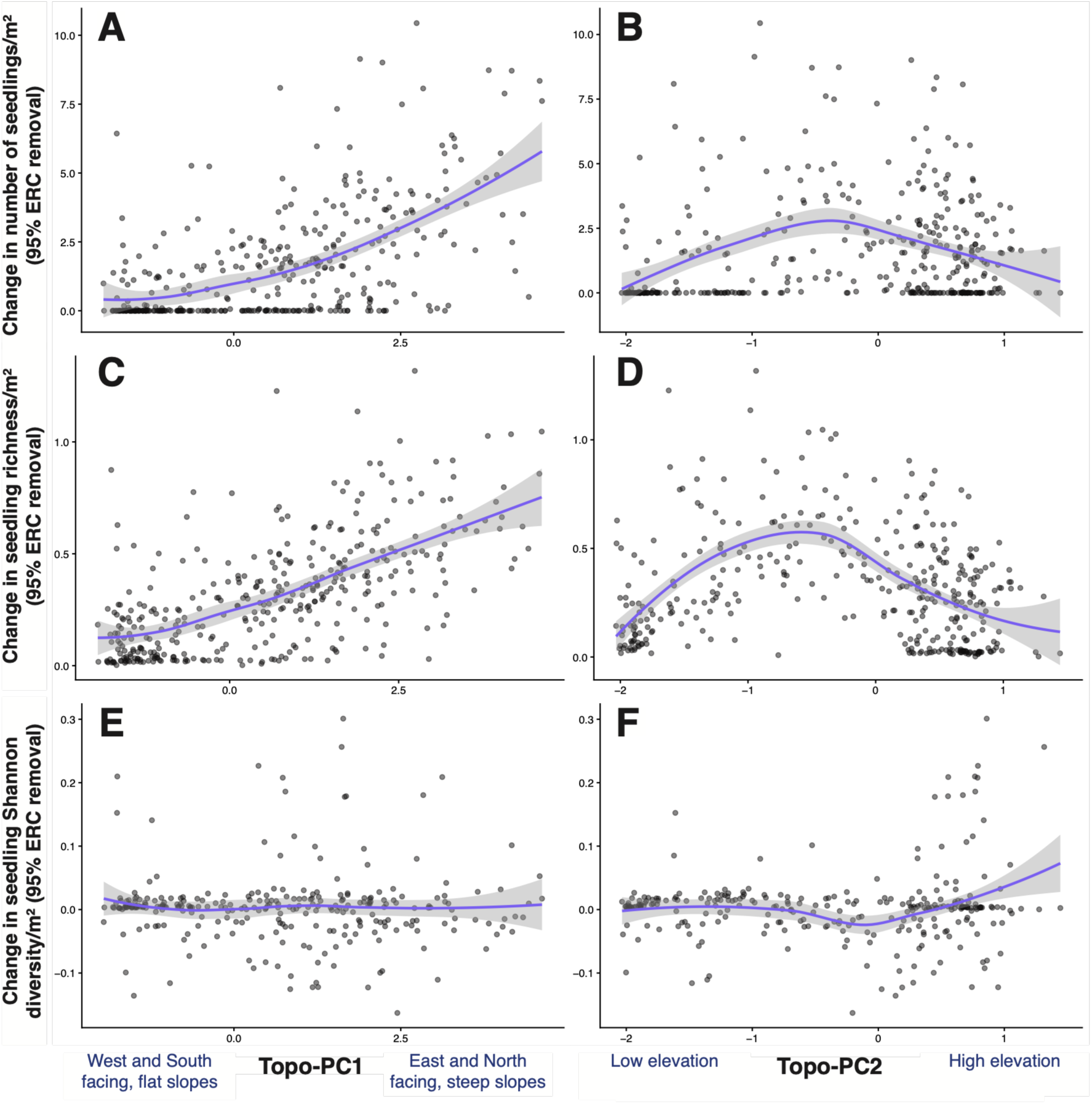
Topographic habitats where simulated ERC managment is predicted to have the greatest understory impact. The predicted change in woody seedling abundance and richness following 95% ERC removal across a topographic gradient. (A) Change in seedling abundance per m² along Topo-PC1 (aspect and slope), (B) change in seedling abundance along Topo-PC2 (elevation), (C) change in seedling richness along Topo-PC1, (D) change in seedling richness along Topo-PC2, (E) change in seedling Shannon diversity along Topo-PC1, and (F) change in seedling Shannon diversity along Topo-PC2.

Simulated ERC removal was predicted to have little effect on woody seedling community composition, exhibiting only modest shifts with 95% ERC basal area removal (Figure 6A). However, Continued ERC encroachment was predicted to cause shifts in woody seedling communities in management scenarios where 10% of the basal area of American basswood, green ash, or common hackberry were replaced by ERC (Figure 6B). The community shift resulting from replacement was directional and significant for bur oak (*p* = 0.007), American basswood (*p* < 0.001), green ash (*p* < 0.001), and common hackberry (*p* < 0.001). The woody species predicted to be most associated with a shift to ERC were the shrubs *Prunus americana* Marsh*., Prunus virginiana* L.*, Rhus glabra* L.*, Rosa arkansana* Porter*, Rubus occidentalis* L., and *Symphoricarpos occidentalis* Hook. (Table S7). The species predicted to be least associated with a shift to ERC were climbing vines *Celastrus scandens* L. *and Lonicera dioica L.,* shrub *Ribes americanum* Mill., and tree seedlings *Celtis occidentalis, Fraxinus pennsylvanica, Ostrya virginiana* (Mill.) K. Koch*, Pinus ponderosa* Douglas ex C. Lawson, and *Tilia americana*.

**Figure 6:**
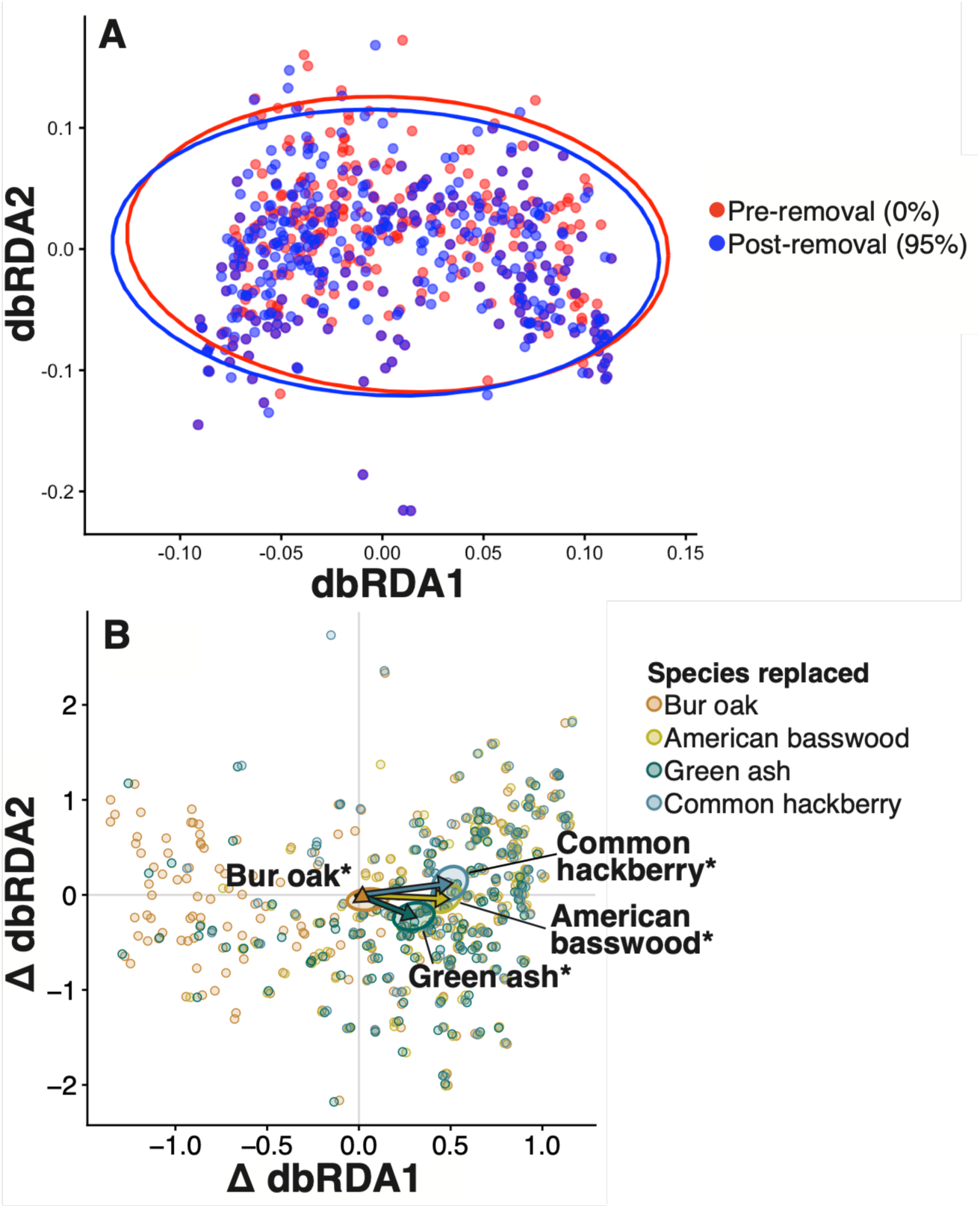
Predicted change in woody seedling community under simulated ERC management and encroachment scenarios. (A) Distance-based redundancy analysis (dbRDA) ordination of projected difference in seedling community composition before and after simulated removal of 95% of ERC. (B) dbRDA ordination of predicted shifts in woody seedling communities under encroachment scenarios following replacement of 10% of a focal species abundance with ERC. Arrows originate at the mean observed community and end at the mean predicted community. The arrow magnitude indicates the expected seedling community response to ERC replacement per species. Points represent communities with 10% ERC replacement of a focal species. Ellipses are 95% bootstrap confidence regions. Stars indicate compositional shifts are significant and directional.

## 4. Discussion

Although encroachment by ERC is widespread across the Great Plains (Meneguzzo & Liknes, 2015), its effects on woodlands have primarily been assessed in the southern portion of the region (Fowler & Carden, 2024; Fuhlendorf et al., 1997; Hoff et al., 2018; van Els et al., 2010; Yager & Smeins, 1999). Whether woodland communities in the ecologically and biogeographically distinct Northern Great Plains respond similarly to ERC encroachment has been an open question. Our study helps fill this knowledge gap using a neighborhood modeling approach comparing the strength of ERC’s effects relative to other co-dominant canopy tree species — an approach that has not previously been used to quantify the impacts of ERC encroachment in woodlands. ERC, compared to other focal species, uniquely suppressed woody seedling recruitment, as ERC dominance was strongly associated with reduced woody seedling abundance, richness, and Shannon diversity, and increasing dominance of other canopy species exhibited marginally negative to positive effects. These differences in recruitment were also exhibited by strong shifts in the species composition of woody seedlings, which differed between understories where ERC versus other species dominated the canopy, aligning with results from the Southern Great Plains (van Els et al., 2010).

Our findings suggest that ERC affects woody seedling communities primarily through shading effects. ERC disproportionately reduced understory light availability compared to other canopy species in low-light environments, which in the Niobrara woodlands are canyon and upland forest habitats. These habitats support more diverse understory plant communities, compared to other topographic habitats (McNichol et al. 2026). Similar reductions in understory light have been reported for ERC-invaded woodlands in the Southern Great Plains (Fuhlendorf et al., 1997; van Els et al., 2010) suggesting that light limitation may be a consistent mechanism in forests and woodlands impacted by ERC across the region. In contrast, we found little evidence that ERC alters soil properties at our site, despite sometimes inconsistent reports of changes in soil pH, organic matter, and C storage elsewhere (Arend, 1948; Broadfoot, 1951; Norris et al., 2001). Regional variation in soils, topography, and climate may be the cause of the variability in the reported belowground effects of ERC encroachment.

Identifying where ERC impacts the understory the most and where management investments will yield the greatest conservation returns is critical for woodland conservation. While simulations of ERC management using our empirical models have disadvantages, experimental and field studies of management strategies do as well, owing to the long timescales of forest recovery. If we cannot tell whether management is working until years later, we may discover failure only after the opportunity for restoration has passed. Our simulations provide indications of how these woodlands would respond to ERC thinning and further encroachment that can serve as decision tools for manager. We found that the "defend the core" approach would yield the largest absolute increases in woody seedling abundance and species richness, whereas ERC removal from heavily encroached areas would yield the greatest proportional gains. ERC thinning in habitats with the highest levels of understory diversity yield the greatest conservation impacts. These findings underscore the importance of coupling ecological research with management planning (Carter et al., 2020; Cravens et al., 2026) to strategically identify where and how much ERC thinning may yield the greatest return on the investment of scarce conservation resources. Our approach provides a framework for identifying management strategies that are tailored to local ecological conditions rather than relying on generalized prescriptions.

### 4.1. ERC as a woody encroacher

Woody encroachment was originally conceived as population expansion of woody species into open habitats, such as grasslands and savannas, in a process resembling biome conversion (S. Archer et al., 1988; S. R. Archer et al., 2017; Ding & Eldridge, 2024). ERC is a well-known woody encroacher in grasslands in the Great Plains, and our results show that its increasing dominance in treed ecosystems can also be a form of woody encroachment. This process also resembles mesophication in forests, in which altered fire regimes produce denser canopies and compositional change (Nowacki & Abrams, 2008).

Common hackberry, which land managers at our site consider a driver of mesophication, is a useful comparison. Common hackberry, a shade-tolerant, highly fecund mesic species, has flourished in the shaded understory of forests absent of fire (Abrams & Napp, 1986; Ortmann & Contreras, 2020). At our site, both common hackberry and ERC negatively affected woody seedling diversity. Like common hackberry, ERC is one of several fire-intolerant, shade-casting species that that can rapidly increase in density when fire is excluded, leading to changes in the forest understory.

Overdominance, even of native species, can alter the understory and, over long term, result in forest community shifts. In central hardwood forests, overdominance of American beech (*Fagus grandifolia* Ehrh.; Fagaceae) is associated with denser canopy cover and a lack of oak recruitment, effects that are predicted to make forests less climate-resilient (Palus et al., 2018). Similarly, overdominance of pawpaw (*Asimina triloba* (L.) Dunal; Annonaceae) in a temperate forest in the central USA was associated with reduced local species diversity, community size, and beta-diversity (Wassel & Myers, 2025). Seedling suppression suggests that ERC-dominated stands are on a trajectory toward lower woody diversity and a canopy increasingly composed of ERC.

Of the mechanisms we tested regarding how ERC affects seedling communities, light limitation was the best supported. ERC reduced understory PPFD more than the other focal species, particularly where light was already low, suggesting that severe light limitation caused by ERC canopy densification is a mechanism that inhibits vegetation growth in the understory. This is consistent with the oak recruitment decline related to light availability reported in other ERC-encroached woodlands (Dey et al., 2014; Schneider et al., 2011). Conversely, ERC was not associated with drier soils, altered soil pH, or changes in soil organic carbon, suggesting that water limitation, leaf-litter acidification, and altered decomposition are unlikely drivers at our site, though the sandy, rapidly draining soils may have limited our ability to detect moisture differences. Mechanisms such as allelopathy would be worth further investigation; there is evidence that compounds leached from ERC roots or foliage are allelopathic and inhibit germination of other species (Bennion & Ward, 2023; Smith, 1986).

### 4.2. Management implications

In forests with encroaching ERC, managers face a dilemma: removal is costly, may have unwanted collateral impacts on fragile ecosystems through abrupt changes in light availability or increasing erosion, and monitoring can take years to detect a response. These risks are greatest systems of high conservation value, such as the woodlands in the Niobrara River Valley. Empirical models of understory response to ERC management simulate alternative scenarios with predictions grounded in field data. These predictions are not a replacement for experimental removal and monitoring, and they rest on two assumptions: they represent years of accumulated effects of ERC encroachment, and they assume the “new” level of ERC dominance after management is maintained over time through repeated treatments that prevent additional encroachment. While these predictions are not a replacement for experimental removal and monitoring, they can help prioritize where and how to act first.

Our scenarios suggest that what areas should be prioritized for management depend on the management goal. For maximizing woody seedling abundance and richness recovery after management, our models favored sites with 30–50% pre-removal ERC dominance (Figure 4A, 4C). This is somewhat in line with the “defend the core” strategy, focusing on managing ERC in areas with lower levels of encroachment. However, Shannon diversity did not follow this strategy as clearly. Removing ERC from sites where it made up less than 50% of basal area was predicted to have neutral to negative effects on woody seedling diversity (Figure 4E). Because diversity is correlated non-linearly to ERC whereas other response variable effects were linear, diversity outcomes are more challenging to predict. One explanation for this is that ERC at low abundance adds to structural and functional heterogeneity in this deciduous-dominated woodland. ERC is one of two species of conifer (the other being Ponderosa pine, *Pinus ponderosa*). It is important to point out that the predicted diversity loss at low pre-removal ERC encroachment levels is very modest relative to the range of woody diversity across the Niobrara Plot (Figure S5A), so there may be a warranted tradeoff. Treating areas with low levels of encroachment likely costs less and slows the spread into new areas, as ERC seedlings are dispersed outward from source populations (Donovan et al., 2018; D. T. Fogarty et al., 2022). Delaying treatment may reduce the local seed sources available to reestablish the woody community (Maclean et al., 2018; Wang et al., 2022), and allowing encroachment to continue could cause a much larger but delayed diversity decrease. Overall, our simulations indicate that prioritizing ERC removal in low to moderate areas of encroachment has the most potential to increase woody seedling abundance and richness at a small cost to diversity.

Simulated benefits also varied with topography. Increases in seedling abundance and richness were greatest on steep, north- and east-facing slopes at intermediate elevations (high topo-PC1 and intermediate topo-PC2). At our site, steeper slopes are found in the canyons, which are located at median elevations between the high elevation prairie ecotone and low elevation floodplain. Canyons are cooler, more sheltered, and have distinct microclimates (McNichol et al., 2024). They are generally associated with more diverse and dense plant communities that harbor sensitive species (McNichol & Russo, 2026) and higher adult tree diversity (McNichol et al., 2024). This underscores the need for management prioritization in diverse, sensitive areas.

Previous studies have examined herbaceous “winners” and “losers” of ERC encroachment (Domeier et al., 2025; Limb et al., 2010), but this study is novel in that we examined woody seedlings in a woodland. ERC encroachment is predicted to shift seedling communities toward disturbance-tolerant shrubs adapted to dry, high-light, or disturbed conditions (e.g. *Prunus americana, Rhus glabra, Rosa arkansana*), and away from trees and climbing vines/shrubs with fruits and flowers with high value to wildlife and pollinators (e.g. *Lonicera dioica, Pinus ponderosa, Ribes americana*). For management goals of increasing seedling recruitment or protecting wildlife habitat, these models are particularly useful, and out models indicate that ERC removal protects wildlife habitat.

## 5. Conclusions

Our study in a Northern Great Plains woodland with great conservation importance provides a clear answer to our question, “can a woody encroaching species affect woodlands?” Yes, eastern redcedar dramatically alters the community structure of woodland understory plants. Suppressed recruitment of seedlings, the future of the forest, resulted in reduced seedling abundance, richness, diversity, and altered seedling composition — effects which were likely driven by reductions in understory light. Continued ERC encroachment is likely to have effects that continue beyond the seedling stage, preventing juveniles from reaching the canopy and gradually shifting canopy composition toward even greater ERC dominance. In our study site and elsewhere in the Niobrara River valley, local ERC dominance in woodlands can be as great as 100%. Replacing a diverse community of canopy trees with ERC is expected to cascade to affect birds, mammals, and other groups, likely reducing diversity at higher trophic levels, since homogenization of the vegetation would lead to fewer ecological niches (Chapman et al., 2004; Horncastle et al., 2005), which is associated with reduced ecosystem services (Cardinale et al., 2012; Turnbull et al., 2016) Therefore, our study suggests that early detection and purposeful, empirically-guided management to limit ERC encroachment in woodlands is essential to maintaining woodland diversity and ecosystem services.

## Supporting information

Full supplemental tables and figures

## Acknowledgements

We want to thank The Nature Conservancy and its staff at the Niobrara Valley Preserve for logistical support. We also want to thank members of the Niobrara Forest Plot field crews over the years: Emma Decker, Collin Eaton, Sakia Fields, Jamie Fuqua, Jennifer Foster, Joseph Eastman, Townson Lemansky, Jack Luth, Susana Moyer, Ashley Newman, Cooper Steffen, and Patrick Wilson. Thanks to Chad Brassil, Daniel Gschwentner, Clinton Leach, and Dan Metz for statistical advice.

## CRediT author statement

**Lillian Hoffart:** Conceptualization, Methodology, Formal Analysis, Investigation, Writing - Original Draft. **Bailey H. McNichol:** Methodology, Investigation, Writing - Review & Editing. **Abigail Ridder:** Methodology, Investigation, Writing - Review & Editing. **Brittni McGuire:** Methodology, Investigation, Writing - Review & Editing. **Sabrina E. Russo:** Conceptualization, Methodology, Resources, Writing – Review & Editing, Supervision.

## Declaration of generative AI and AI-assisted technologies in the manuscript preparation process

During the preparation of this work the author(s) used Claude (Anthropic) to revise writing and debug analysis code. After using this tool/service, the author(s) reviewed and edited the content as needed and take(s) full responsibility for the content of the published article.

## Data citation

https://doi.org/10.71964/145

