## Supplementary material for "Can a woody encroaching species affect woodlands? Impacts of eastern redcedar (*Juniperus virginiana*) overdominance on woodland understories and management scenarios": Full supplemental tables and figures

**Table S1: Data collected in the Niobrara Plot to address research questions.** In cases in which data were not collected from all 1 $m^2$ subplots, a subset of subplots was chosen using stratified random sampling to represent the full range of habitat conditions throughout the plot (McNichol et al., 2024).

| **Dataset** | **Research Question Model Variables** | **Time Period of Data Collection** | **Scale of Data Collection** |
|---|---|---|---|
| Niobrara Forest Plot inventory and topographic data | Q1-Q3: Predictor variable: Neighborhood dominance, topographic position | Summer 2018 - 2019 | Census of all stems ≥ 1 cm DBH in 20.2 ha plot |
| Woody seedling census | Q1: Response variables: Seedling abundance, Shannon diversity, richness, and evenness | Summers 2021, 2022 | 504 subplots |
| Ground cover survey | Q2: Response variable: Percent vegetative groundcover (woody and herbaceous) | Summer 2022 | 504 subplots |
| Herbaceous plant survey | Q2: Response variable: Herbaceous species richness | July – August 2021 | 78 subplots |
| Soil properties | Q3: Response variables: Soil pH, soil organic matter | August 2022 | Adjacent to 50 subplots |
| Soil moisture | Q3: Response variables: Soil volumetric water content | Averages of data collected monthly (May - October 2021) | Adjacent to 74 subplots |
| Estimated PPFD | Q3: Response variable: Understory light availability | July 2022 | 504 subplots |

**Figure S1: Frequency of the neighborhood dominance of each species on a 0-1 scale.** ERC = eastern redcedar.

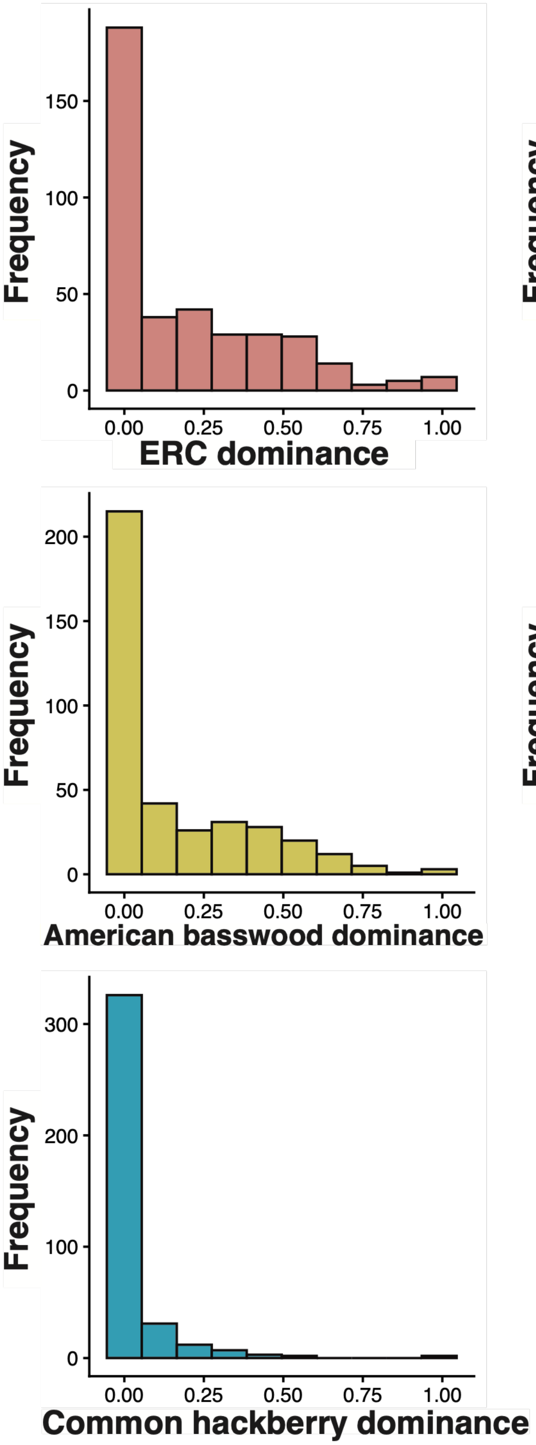

**Table S2: Factor loadings from the principal component analysis (PCA) of topographic variables.**

| | **PCA factor** | |
|---|---|---|
| **Variable** | **1** | **2** |
| Slope | 0.582 | 0.062 |
| Elevation | -0.108 | 0.994 |
| Eastness | 0.568 | 0.067 |
| Northness | 0.571 | 0.059 |

**Table S3: Model selection comparing models with no interactions, 2-way interactions, and 3-way interactions for each response variable.** Bold font indicated the most supported model used in analyses. The 2-way interaction model included interactions each species' dominance and topo-PC1, and the 3-way interaction model included the 2 two-way interactions plus the three-way interactions between each species' dominance × topo-PC1 × topo-PC2. ELPD = expected log pointwise predictive density; SE = standard error; VWC = volumetric water content; LOI = loss on ignition; PPFD = photosynthetic photon flux density.

| | **No interactions** | | **2-way interactions** | | **3-way interactions** | |
|---|---|---|---|---|---|---|
| | **ΔELPD** | **ΔSE** | **ΔELPD** | **ΔSE** | **ΔELPD** | **ΔSE** |
| Woody seedling abundance | **-11.4** | **6.3** | -7.8 | 4.7 | 0.0 | 0.0 |
| Woody seedling richness | **-4.8** | **3.5** | 0.0 | 0.0 | -1.3 | 3.5 |
| Woody seedling evenness | **0.0** | **0.0** | -3.8 | 2.3 | -4.8 | 6.3 |
| Woody seedling diversity | **-9.3** | **6.5** | -0.8 | 4.5 | 0.0 | 0.0 |
| Vegetative ground cover | **-2.9** | **3.3** | 0.0 | 0.0 | -4.7 | 3.4 |
| Herbaceous species richness | **0.0** | **0.0** | -1.5 | 2.1 | -10.4 | 5.5 |
| Soil VWC | **0.0** | **0.0** | -3.4 | 2.8 | -0.3 | 5.6 |
| Soil pH | **0.0** | **0.0** | -6.1 | 1.7 | -12.6 | 4.6 |
| Soil organic matter (% LOI) | **0.0** | **0.0** | -3.8 | 2.1 | -17.9 | 4.3 |
| PPFD | **0.0** | **0.0** | -39.9 | 10.9 | -38.6 | 14.8 |

**Table S4: Spatial autocorrelation assessment for each model**. For each model response variable, the following is listed: Moran's *I* for the model without accounting for spatial autocorrelation, whether a conditional autoregressive (CAR) term was included, and the Moran's *I* values after the addition of the component to the model. Woody seedling data was collected in years, so each year was assessed for spatial autocorrelation separately. * Models did not converge when adding a CAR term.

| **Response Variable** | **Moran's *I* without accounting for spatial autocorrelation** | **CAR term?** | **Moran's *I* including the CAR term** |
|---|---|---|---|
| Woody seedling abundance | 2021: 0.132, $p < 0.001$<br>2022: 0.130, $p < 0.001$ | Yes | 2021: 0.016, $p = 0.228$<br>2022: 0.022, $p = 0.155$ |
| Woody seedling species richness | 2021: 0.044, $p = 0.044$<br>2022: -0.005, $p = 0.519$ | Yes | 2021: 0.005, $p = 0.374$<br>2022: -0.042, $p = 0.912$ |
| Woody seedling evenness | 2021: 0.030, $p = 0.157$<br>2022: 0.027, $p = 0.175$ | No | - |
| Woody seedling Shannon diversity | 2021: 0.192, $p < 0.001$<br>2022: 0.192, $p < 0.001$ | No | * |
| Vegetative groundcover (%) | 0.162, $p < 0.001$ | Yes | -0.070, $p = 0.996$ |
| Herbaceous species richness | 0.027, $p = 0.210$ | No | - |
| Soil VWC (%) | 0.108, $p = 0.006$ | No | * |
| Soil pH | -0.029, $p = 0.543$ | No | - |
| Soil organic matter (% LOI) | 0.071, $p = 0.064$ | No | - |
| PPFD | 0.319, $p < 0.001$ | No | * |

**Table S5: Mean and 95% credible intervals for slope parameters estimating the effects of each focal species' dominance on the understory plant community.** Bold font indicates slope parameters with 95% credible intervals excluding zero. ERC = eastern redcedar; VWC = volumetric water content; LOI = loss on ignition; PPFD = photosynthetic photon flux density. *Because of the quadratic term, the slope and 95% credible intervals for these species in the woody seedling diversity model were determined at a dominance of 0.50.

| Response variable | Species | Slope (95% credible interval) |
|---|---|---|
| Woody seedling abundance | ERC | **-1.693 (-2.598, -0.804)** |
| | Bur oak | -0.710 (-1.451, 0.008) |
| | American basswood | 0.288 (-0.818, 1.406) |
| | Green ash | -1.138 (-2.325, 0.011) |
| | Common hackberry | 0.150 (-1.524, 1.814) |
| Woody seedling species richness | ERC | **-0.655 (-1.063, -0.250)** |
| | Bur oak | 0.033 (-0.297, 0.365) |
| | American basswood | 0.337 (-0.099, 0.770) |
| | Green ash | 0.040 (-0.455, 0.532) |
| | Common hackberry | -0.579 (-1.350, 0.164) |
| Woody seedling evenness | ERC | 0.461 (-0.335, 1.250) |
| | Bur oak | **0.669 (0.044, 1.261)** |
| | American basswood | 0.201 (-0.520, 0.904) |
| | Green ash | 0.816 (-0.011, 1.633) |
| | Common hackberry | -0.585 (-1.980, 0.885) |
| Woody seedling diversity | ERC * | **-1.220 (-1.979, -0.439)** |
| | Bur oak * | -0.195 (-0.446, 0.058) |
| | American basswood | -0.307 (-0.665, 0.047) |
| | Green ash * | 0.121 (-0.171, 0.408) |
| | Common hackberry | **-0.701 (-1.249, -0.138)** |
| Vegetative ground cover | ERC | -0.290 (-0.845, 0.266) |
| | Bur oak | -0.084 (-0.522, 0.352) |
| | American basswood | 0.386 (-0.280, 1.055) |
| | Green ash | 0.006 (-0.671, 0.689) |
| | Common hackberry | 0.172 (-0.837, 1.205) |
| Herbaceous species richness | ERC | -0.056 (-0.622, 0.515) |
| | Bur oak | 0.059 (-0.392, 0.504) |
| | American basswood | 0.137 (-0.447, 0.706) |
| | Green ash | 0.502 (-0.183, 1.185) |
| | Common hackberry | 0.677 (-0.895, 2.302) |
| Soil VWC | ERC | 0.258 (-0.370, 0.891) |
| | Bur oak | -0.306 (-0.842, 0.237) |
| | American basswood | 0.190 (-0.545, 0.947) |
| | Green ash | 0.528 (-0.183, 1.270) |
| | Common hackberry | -0.726 (-2.553, 1.075) |
| Soil pH | ERC | -0.950 (-2.056, 0.206) |
| | Bur oak | -0.689 (-1.629, 0.252) |
| | American basswood | -0.129 (-1.204, 0.973) |
| | Green ash | -0.120 (-1.292, 1.069) |
| | Common hackberry | -0.020 (-2.316, 2.226) |
| Soil organic matter (% LOI) | ERC | -0.005 (-0.077, 0.068) |
| | Bur oak | -0.008 (-0.068, 0.054) |
| | American basswood | 0.021 (-0.047, 0.093) |

| | | |
|---|---|---|
| | Green ash | 0.004 (-0.068, 0.077) |
| | Common hackberry | -0.023 (-0.165, 0.121) |
| PPFD | High-light ERC | 0.007 (-0.713, 0.739) |
| | Low-light ERC | **-1.149 (-1.779, -0.517)** |
| | High-light all other species | -0.023 (-0.648, 0.565) |
| | Low-light all other species | -0.0877 (-0.611, 0.437) |

**Figure S2: The relationship between woody seedling abundance and canopy species dominance for five focal species:** (A) eastern redcedar, (B) bur oak, (C) American basswood, (D) green ash, and (E) common hackberry. Solid lines represent the model-predicted relationship between the canopy species and seedling abundance, with shaded ribbons representing 95% credible intervals. All other predictors were held at their mean values. Bold points with error bars show binned means ± 95% confidence intervals, calculated by dividing each species' impact range into 5 equal-width bins finding the average of each bin. The dotted line represents the seedling abundance global mean. (F) The slope estimates and 95% credible interval for each species. Stars indicate credible intervals that exclude zero.

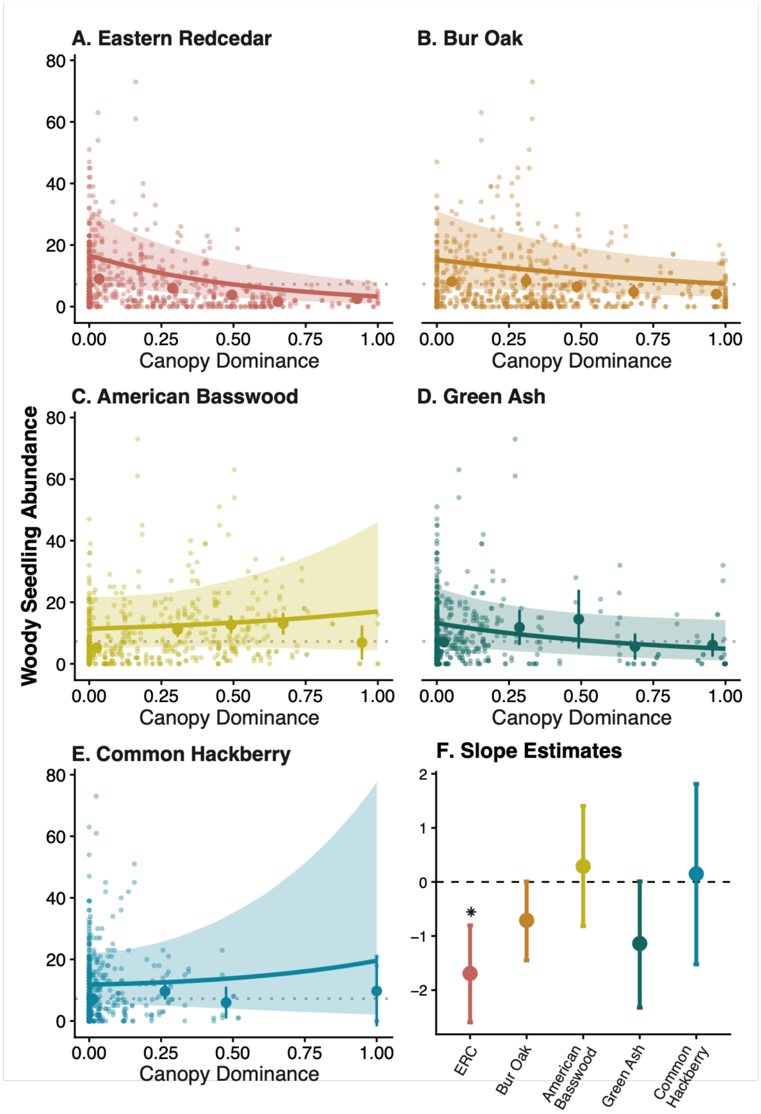

**Table S6: Planned contrasts of the magnitude of eastern redcedar (ERC) dominance versus other focal tree species on the understory.** Positive contrasts indicate that the focal species had a more positive effect than ERC, whereas negative contrasts indicate that the focal species had a more negative effect than ERC. Bold font indicates contrasts with 95% credible intervals excluding zero. VWC = volumetric water content; LOI = loss on ignition; PPFD = photosynthetic photon flux density. *Because of the quadratic term, the slope and 95% credible intervals for these species in the woody seedling diversity model were determined at a dominance of 0.50.

| Model | Focal species vs. ERC | Difference |
|---|---|---|
| Woody seedling abundance | Bur oak | **0.982 (0.138, 1.821)** |
| | American basswood | **1.981 (0.882, 3.078)** |
| | Green ash | 0.554 (-0.645, 1.740) |
| | Common hackberry | **1.843 (0.134, 3.530)** |
| Woody seedling species richness | Bur oak | **0.688 (0.286, 1.091)** |
| | American basswood | **0.992 (0.534, 1.445)** |
| | Green ash | **0.695 (0.167, 1.221)** |
| | Common hackberry | 0.076 (-0.712, 0.838) |
| Woody seedling evenness | Bur oak | 0.209 (-0.615, 1.032) |
| | American basswood | -0.259 (-1.069, 0.559) |
| | Green ash | 0.355 (-0.600, 1.325) |
| | Common hackberry | -1.045 (-2.445, 0.415) |
| Woody seedling diversity | Bur oak * | **1.025 (0.249, 1.786)** |
| | American basswood | **1.341 (0.561, 2.114)** |
| | Green ash * | **0.913 (0.092, 1.717)** |
| | Common hackberry | 0.519 (-0.428, 1.478) |
| Vegetative ground cover | Bur oak | 0.206 (-0.310, 0.723) |
| | American basswood | 0.676 (-0.001, 1.365) |
| | Green ash | 0.297 (-0.404, 0.997) |
| | Common hackberry | 0.462 (-0.565, 1.511) |
| Herbaceous species richness | Bur oak | **0.688 (0.286, 1.091)** |
| | American basswood | **0.992 (0.534, 1.445)** |
| | Green ash | **0.695 (0.1667, 1.221)** |
| | Common hackberry | 0.076 (-0.712, 0.838) |
| Soil VWC | Bur oak | -0.564 (-1.216, 0.104) |
| | American basswood | -0.068 (-0.839, 0.706) |
| | Green ash | 0.270 (-0.484, 1.024) |
| | Common hackberry | -0.983 (-2.833, 0.832) |
| Soil pH | Bur oak | 0.262 (-0.682, 1.189) |
| | American basswood | 0.821 (-0.369, 2.021) |
| | Green ash | 0.830 (-0.421, 2.062) |
| | Common hackberry | 0.930 (-1.468, 3.294) |
| Soil organic matter (% LOI) | Bur oak | -0.004 (-0.065, 0.059) |
| | American basswood | 0.025 (-0.052, 0.105) |
| | Green ash | 0.009 (-0.067, 0.087) |
| | Common hackberry | -0.019 (-0.170, 0.128) |
| PPFD | High-light individuals | -0.030 (-0.691, 0.583) |
| | Low-light individuals | **1.061 (0.468, 1.654)** |

**Figure S3: The relationship between woody seedling species richness and canopy species dominance for five focal species:** (A) eastern redcedar, (B) bur oak, (C) American basswood, (D) green ash, and (E) common hackberry. Solid lines represent the model-predicted relationship between the canopy species and seedling richness, with shaded ribbons representing 95% credible intervals. All other predictors were held at their mean values. Bold points with error bars show binned means ± 95% confidence intervals, calculated by dividing each species' impact range into 5 equal-width bins finding the average of each bin. The dotted line represents the seedling richness global mean. (F) The slope estimates and 95% credible interval for each species. Stars indicate credible intervals that exclude zero.

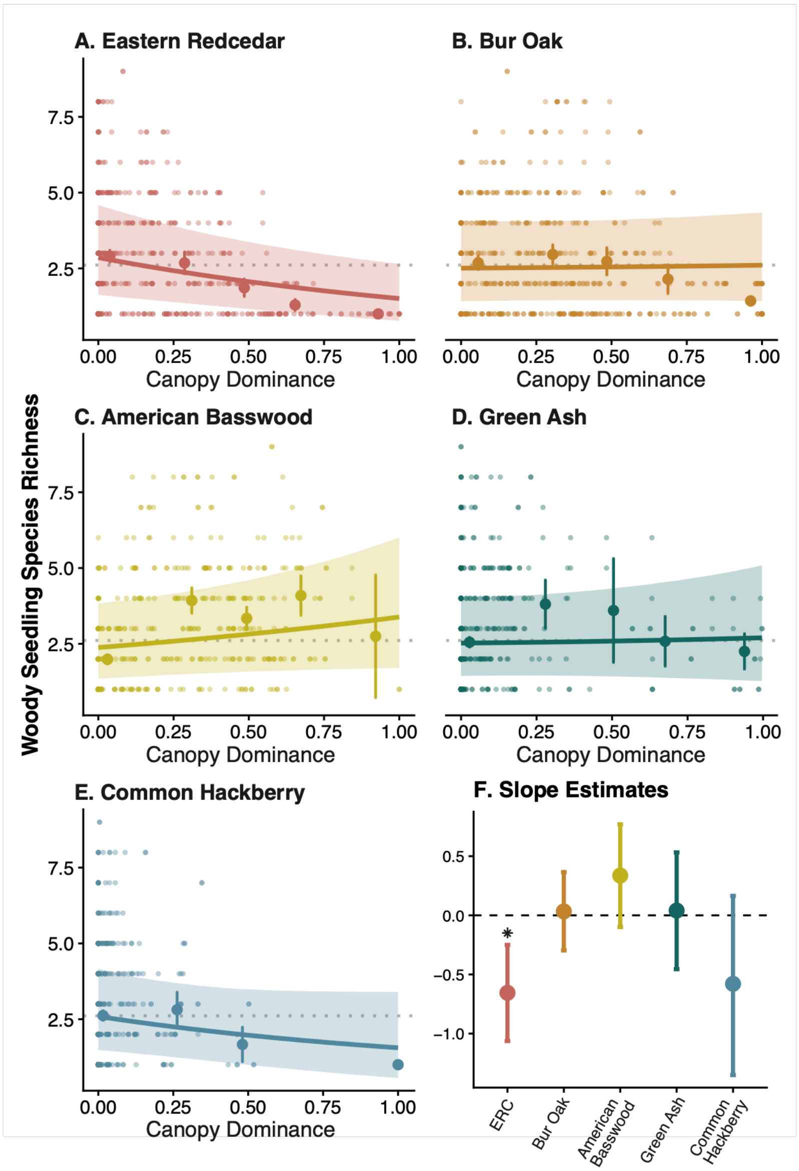

**Figure S4: The relationship between woody seedling evenness and canopy species dominance for five focal species:** (A) eastern redcedar, (B) bur oak, (C) American basswood, (D) green ash, and (E) common hackberry. Solid lines represent the model-predicted relationship between the canopy species and seedling evenness, with shaded ribbons representing 95% credible intervals. All other predictors were held at their mean values. Bold points with error bars show binned means ± 95% confidence intervals, calculated by dividing each species' impact range into 5 equal-width bins finding the average of each bin. The dotted line represents the seedling evenness global mean. (F) The slope estimates and 95% credible interval for each species. Stars indicate credible intervals that exclude zero.

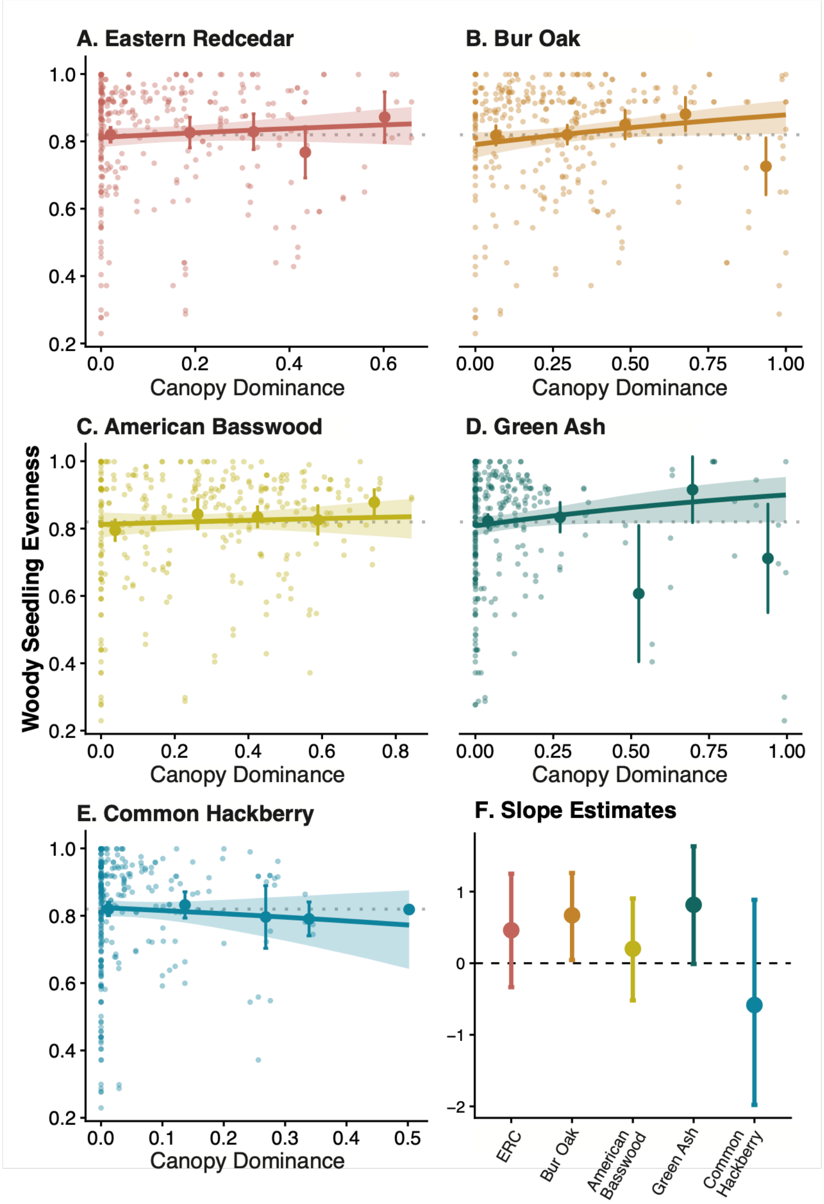

**Figure S5: The relationship between woody seedling Shannon diversity and canopy species dominance for five focal species:** (A) eastern redcedar, (B) bur oak, (C) American basswood, (D) green ash, and (E) common hackberry. Solid lines represent the model-predicted relationship between the canopy species and seedling Shannon diversity, with shaded ribbons representing 95% credible intervals. All other predictors were held at their mean values. Bold points with error bars show binned means ± 95% confidence intervals, calculated by dividing each species' impact range into 5 equal-width bins finding the average of each bin. The horizontal dotted line represents the Shannon diversity global mean. The vertical dashed line represents the dominance where, due to a quadratic term, ERC, bur oak, and green ash slopes were evaluated. (F) The slope estimates and 95% credible interval for each species. Stars indicate credible intervals that exclude zero.

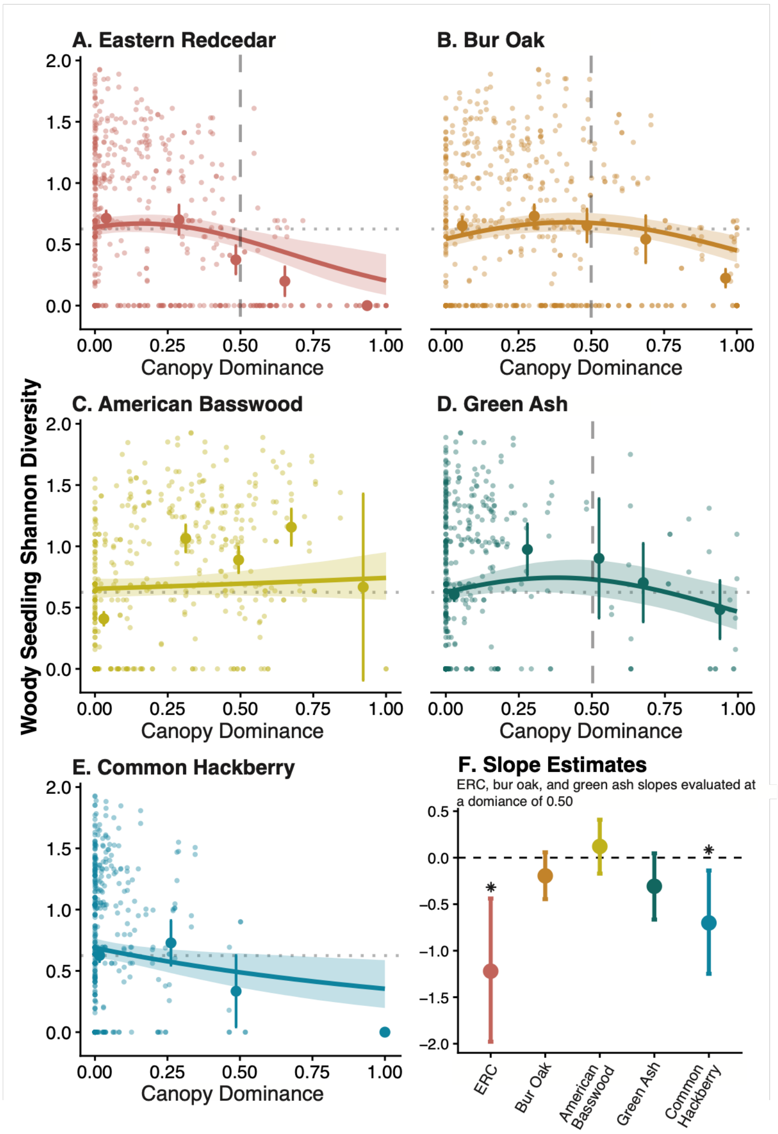

**Figure S6: Variation in understory plant species composition with respect to canopy species dominance and environmental conditions.** Biplot of distance-based redundancy analysis (dbRDA) showing the relationship between focal canopy species dominance (Eastern redcedar (ERC), bur oak, American basswood, green ash, common hackberry), topographic principal components (PC), and understory light (PPFD) on (A) woody seedling community composition, and (B) herbaceous plant community composition. Topography PC1 was associated with more north and easterly facing, steeper slopes, and topography PC2 was associated with higher elevation (see Table S3 for loadings). The length of the vectors approximates the strength of the effect. Percent of variance accounted for by the first two RDA axes is indicated in parentheses. Arrows pointing in similar directions are associated with similar variation in community composition.

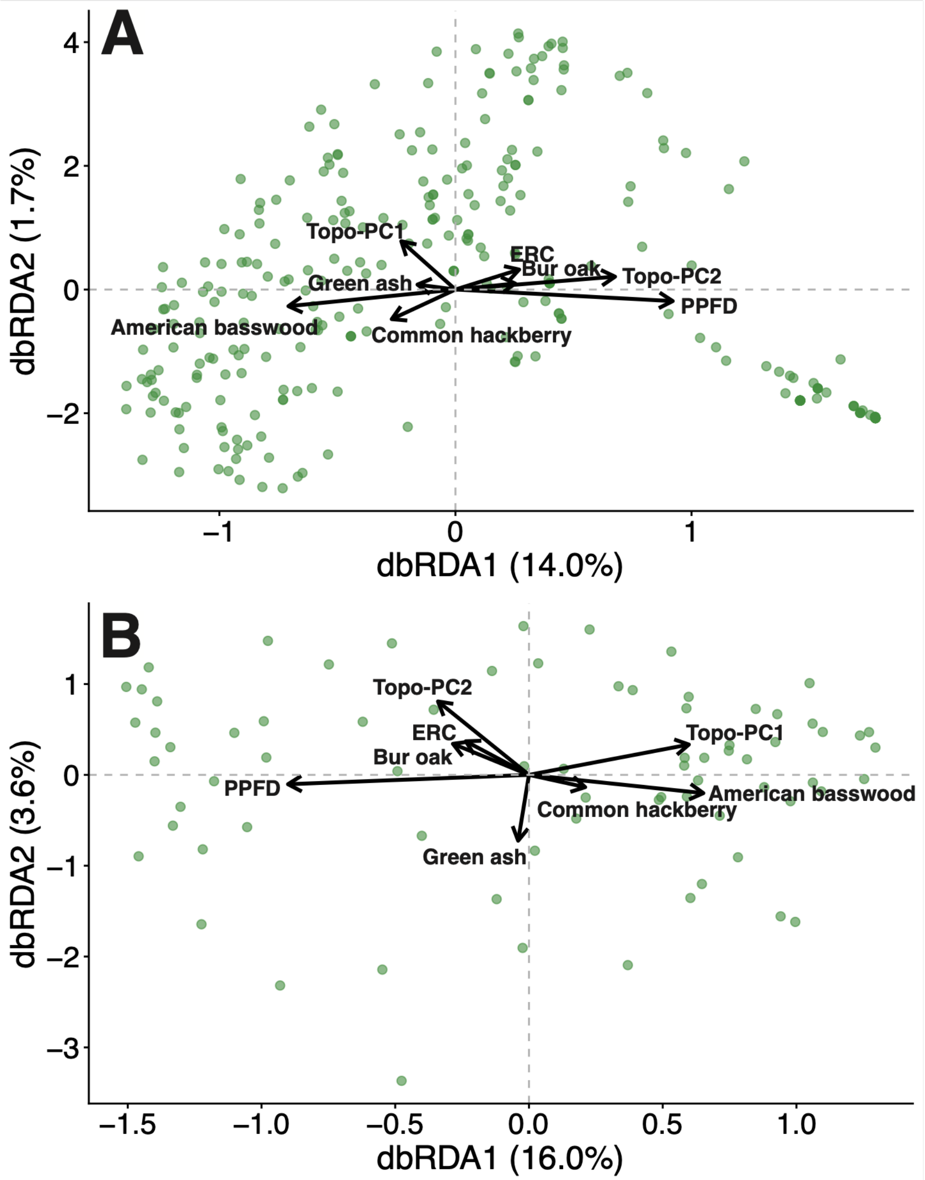

**Table S7: Predicted shifts in understory plant community composition with continued encroachment by eastern redcedar (ERC).** The top five woody species with the greatest and least association with a simulated replacement of 10% of the basal area of a non-ERC focal species with ERC. Negative (orange) values indicate that the understory species will increase with ERC replacement. Positive (blue) values indicate that the understory species will decrease with ERC replacement. Blank cells indicate that woody species was not in the top five species with the greatest and least association with a focal species.

| **Species** | **Bur oak** | **American basswood** | **Green ash** | **Common hackberry** |
|---|---|---|---|---|
| *Celastrus scandens* | -0.681 | | -0.596 | |
| *Celtis occidentalis* | | -0.645 | | -0.726 |
| *Fraxinus pennsylvanica* | | -0.666 | | -0.676 |
| *Lonicera dioica* | -1.275 | -0.894 | -1.244 | |
| *Ostrya virginiana* | -0.506 | -0.655 | -0.561 | -0.618 |
| *Pinus ponderosa* | -0.937 | | -0.900 | |
| *Prunus americana* | | 0.621 | 0.421 | 0.653 |
| *Prunus virginiana* | 0.344 | | | 0.350 |
| *Rhus glabra* | 0.798 | 1.009 | 0.880 | 0.942 |
| *Ribes americanum* | -0.937 | | -0.900 | |
| *Rosa arkansana* | 1.546 | 1.496 | 1.601 | 1.216 |
| *Rubus occidentalis* | | 0.240 | | 0.369 |
| *Smilax lasioneura* | 0.368 | | 0.218 | -0.667 |
| *Symphoricarpos occidentalis* | 0.747 | 0.138 | 0.642 | |
| *Tilia americana* | | -0.696 | | -0.708 |

**Figure S7: The relationship between vegetative ground cover and canopy species dominance for five focal species:** (A) eastern redcedar, (B) bur oak, (C) American basswood, (D) green ash, and (E) common hackberry. Solid lines represent the model-predicted relationship between the canopy species and vegetative ground cover, with shaded ribbons representing 95% credible intervals. All other predictors were held at their mean values. Bold points with error bars show binned means ± 95% confidence intervals, calculated by dividing each species' impact range into 5 equal-width bins finding the average of each bin. The dotted line represents the vegetative ground cover mean. (F) The slope estimates and 95% credible interval for each species. Stars indicate credible intervals that exclude zero.

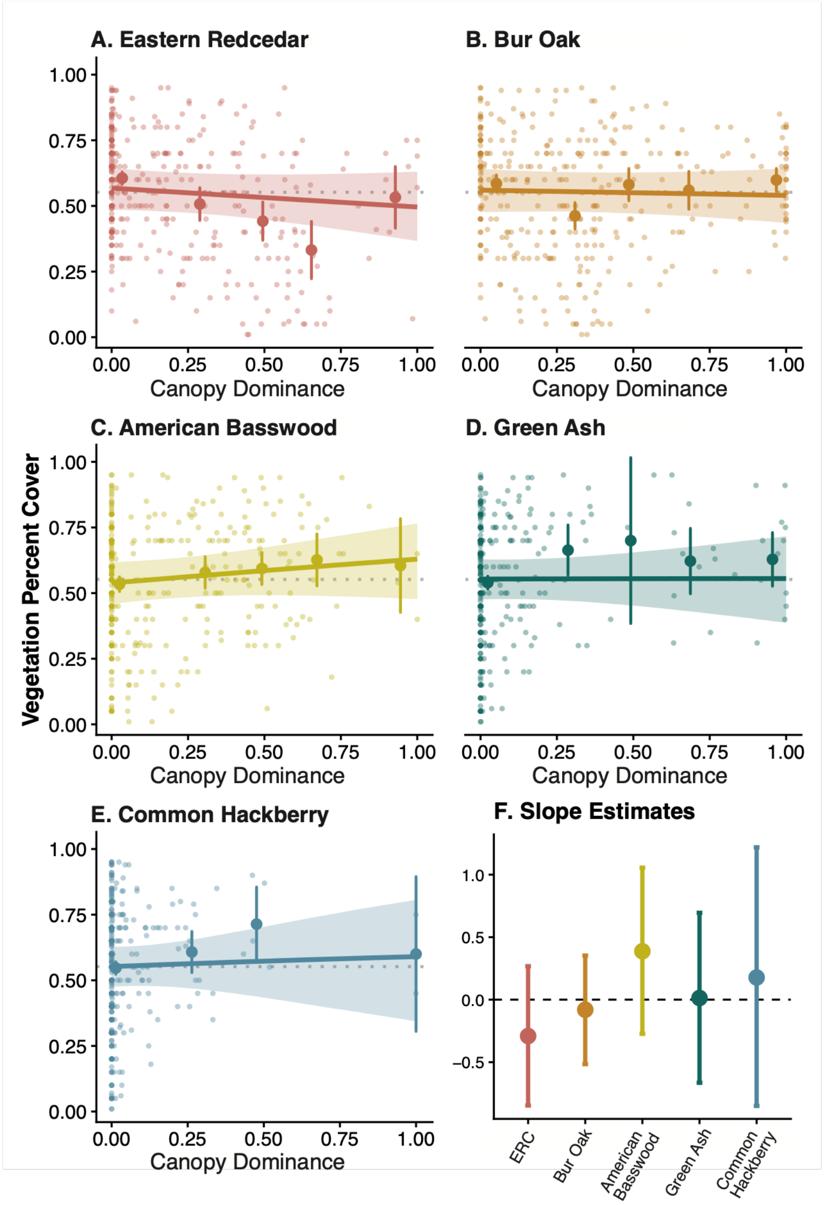

**Figure S8: The relationship between herbaceous species richness and canopy species dominance for five focal species:** (A) eastern redcedar, (B) bur oak, (C) American basswood, (D) green ash, and (E) common hackberry. Solid lines represent the model-predicted relationship between the canopy species and herbaceous richness, with shaded ribbons representing 95% credible intervals. All other predictors were held at their mean values. Bold points with error bars show binned means ± 95% confidence intervals, calculated by dividing each species' impact range into 5 equal-width bins finding the average of each bin. The dotted line represents the herbaceous richness global mean. (F) The slope estimates and 95% credible interval for each species. Stars indicate credible intervals that exclude zero.

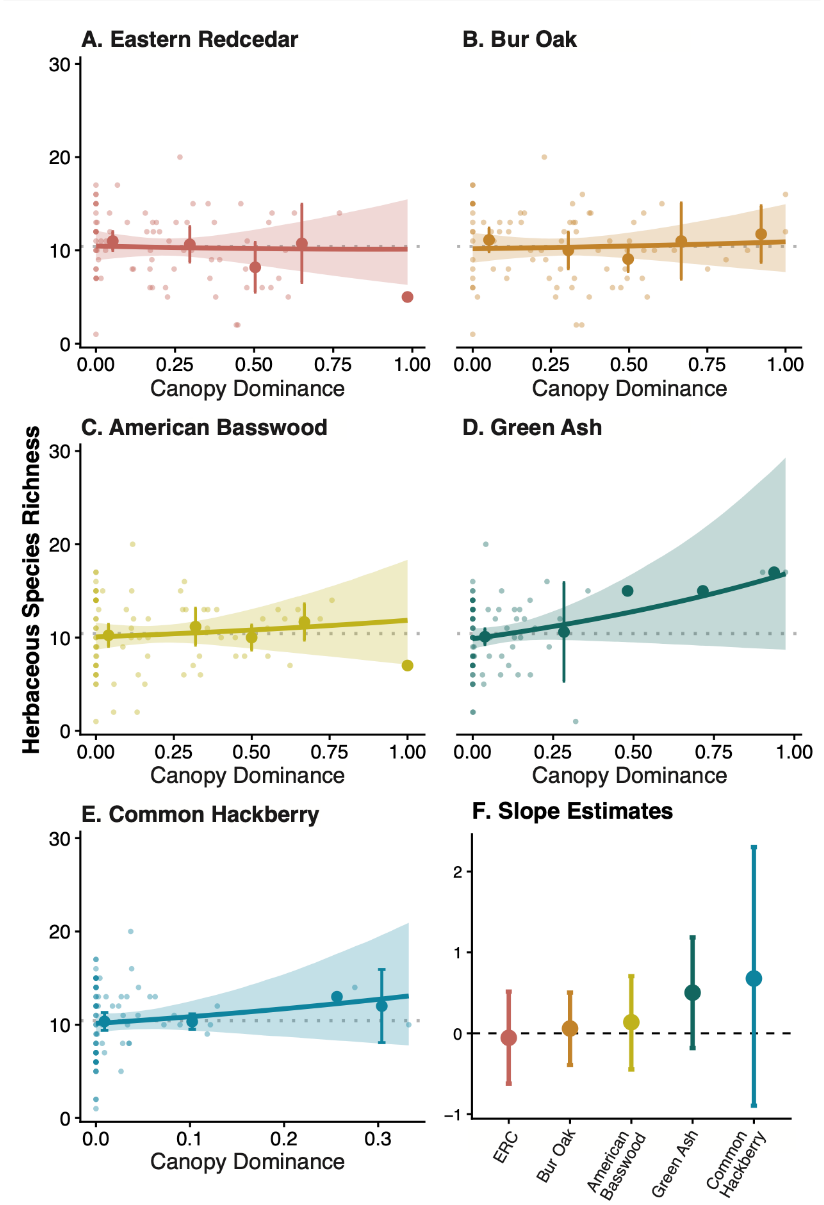

**Figure S9: The relationship between soil volumetric water content (VWC) and canopy species dominance for five focal species:** (A) eastern redcedar, (B) bur oak, (C) American basswood, (D) green ash, and (E) common hackberry. Solid lines represent the model-predicted relationship between the canopy species and soil VWC, with shaded ribbons representing 95% credible intervals. All other predictors were held at their mean values. Bold points with error bars show binned means ± 95% confidence intervals, calculated by dividing each species' impact range into 5 equal-width bins finding the average of each bin. The dotted line represents the soil VWC global mean. (F) The slope estimates and 95% credible interval for each species. Stars indicate credible intervals that exclude zero.

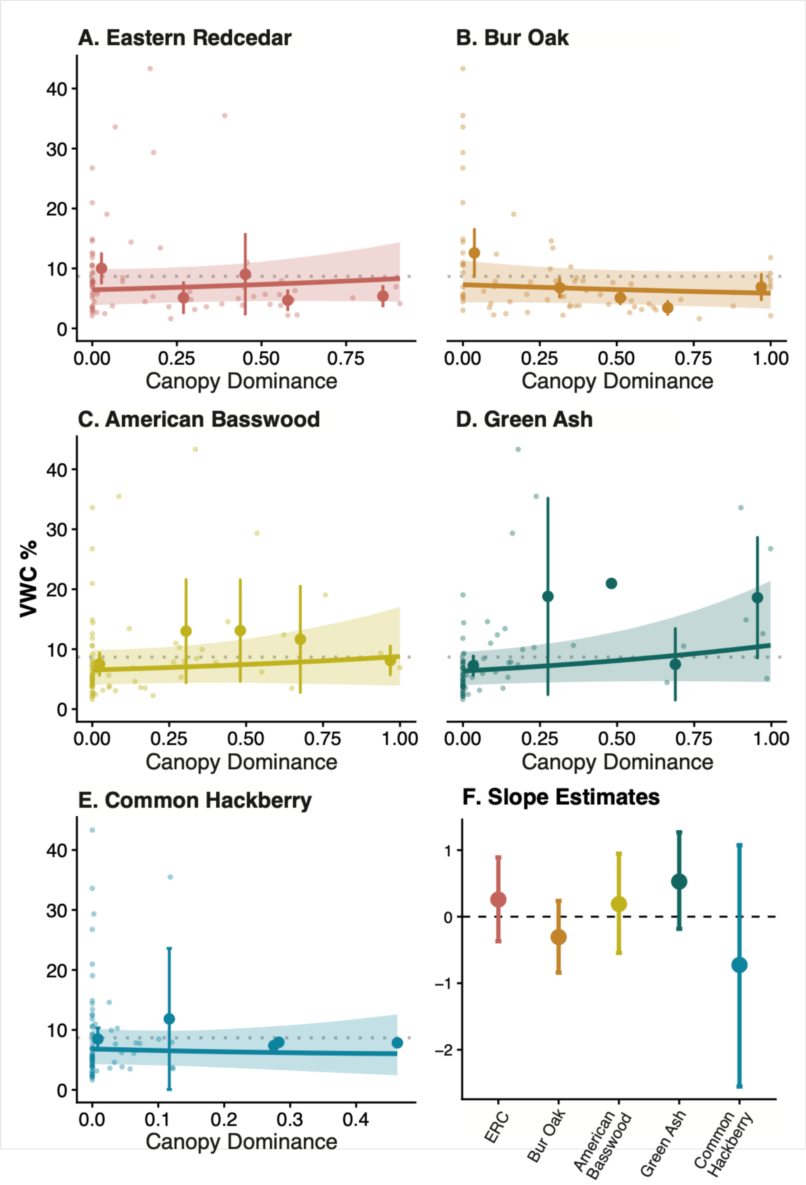

**Figure S10: The relationship between soil pH and canopy species dominance for five focal species:** (A) eastern redcedar, (B) bur oak, (C) American basswood, (D) green ash, and (E) common hackberry. Solid lines represent the model-predicted relationship between the canopy species and soil pH, with shaded ribbons representing 95% credible intervals. All other predictors were held at their mean values. Bold points with error bars show binned means ± 95% confidence intervals, calculated by dividing each species' impact range into 5 equal-width bins finding the average of each bin. The dotted line represents the soil pH global mean. (F) The slope estimates and 95% credible interval for each species. Stars indicate credible intervals that exclude zero.

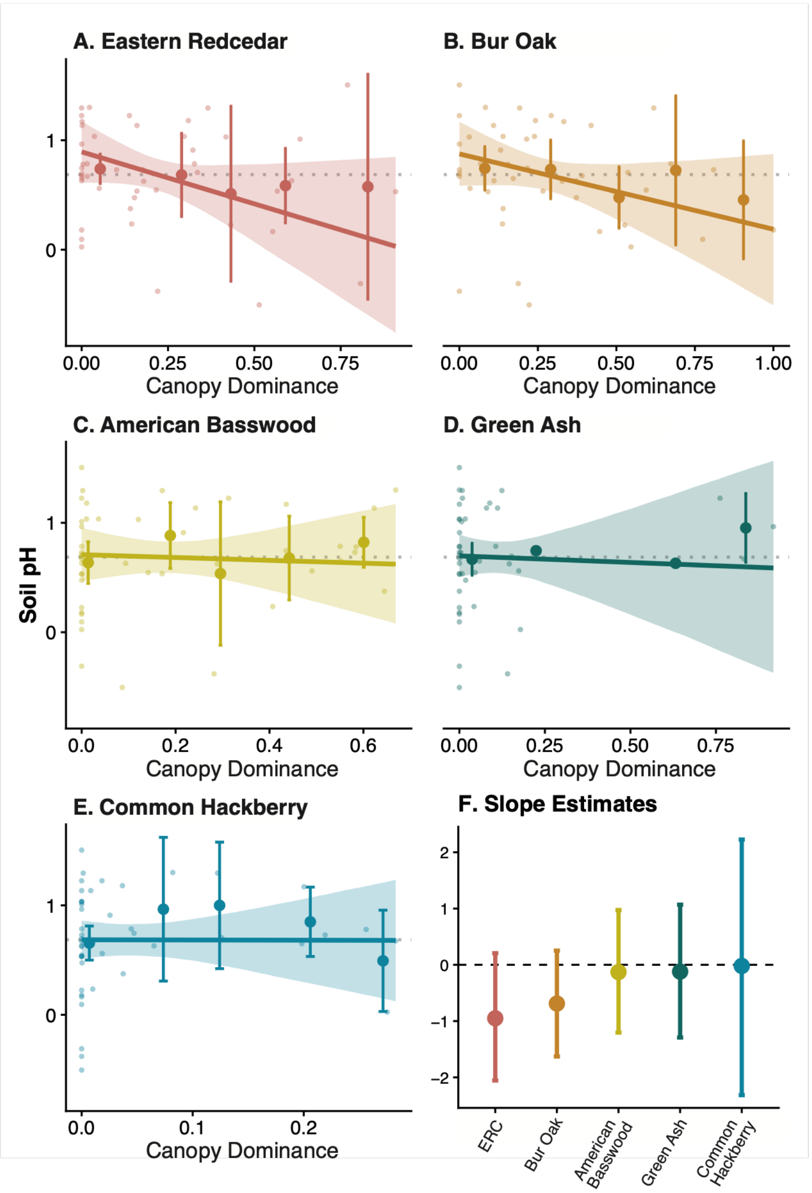

**Figure S11: The relationship between soil organic matter (%LOI) and canopy species dominance for five focal species:** (A) eastern redcedar, (B) bur oak, (C) American basswood, (D) green ash, and (E) common hackberry. Solid lines represent the model-predicted relationship between the canopy species and soil organic matter, with shaded ribbons representing 95% credible intervals. All other predictors were held at their mean values. Bold points with error bars show binned means ± 95% confidence intervals, calculated by dividing each species' impact range into 5 equal-width bins finding the average of each bin. The dotted line represents the soil organic matter global mean. (F) The slope estimates and 95% credible interval for each species. Stars indicate credible intervals that exclude zero.

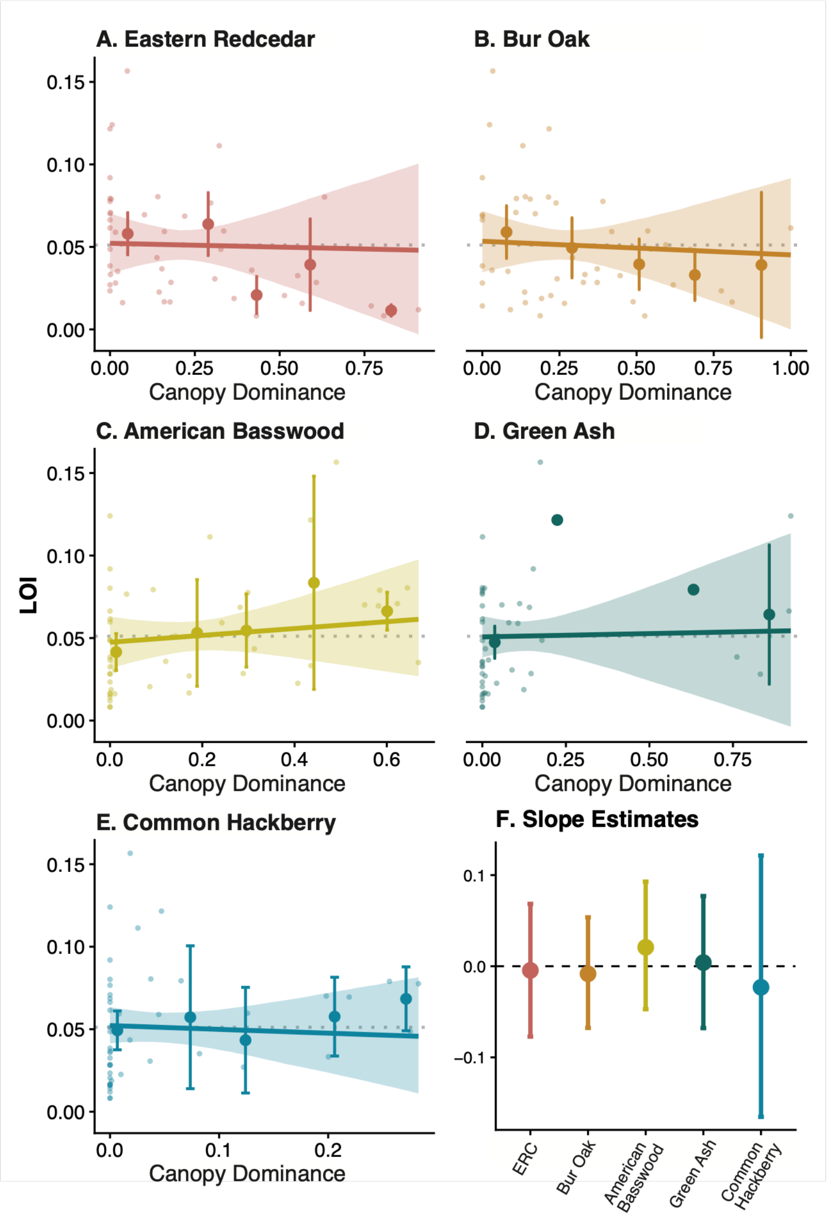

**Figure S12: The relationship between PPFD and canopy species dominance for (A) ERC and (B) four other canopy species** (bur oak, American basswood, green ash, and common hackberry) across different light environments. Solid lines represent the model-predicted relationship between the canopy species and PPFD, with shaded ribbons representing 95% credible intervals. All other predictors were held at their mean values. Bold points with error bars show binned means ± 95% confidence intervals, calculated by dividing each species' impact range into 5 equal-width bins finding the average of each bin. The dotted line represents the soil organic matter global mean. (C) The slope estimates and 95% credible interval for each species. Stars indicate credible intervals that exclude zero.

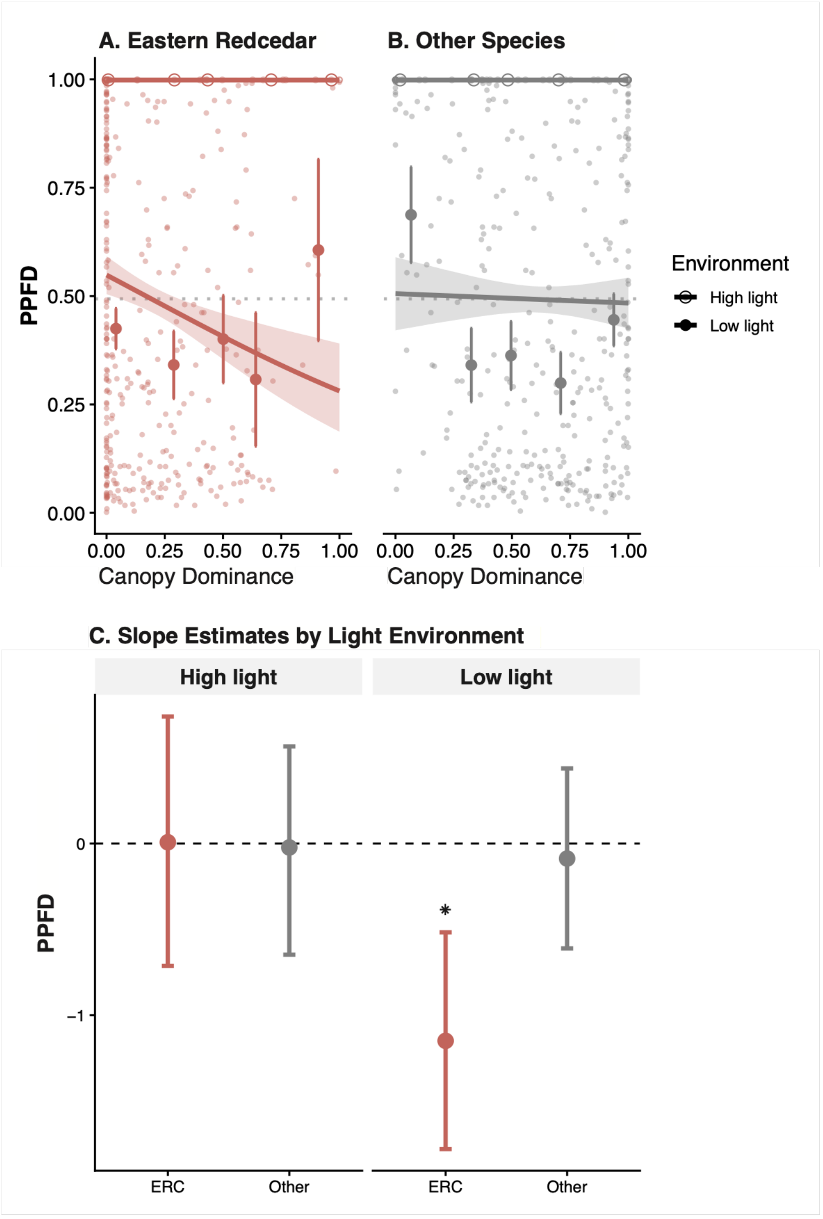
